# Induced Arp2/3 complex depletion increases FMNL2/3 formin expression and filopodia formation

**DOI:** 10.1101/2020.12.03.409573

**Authors:** Vanessa Dimchev, Ines Lahmann, Stefan A. Koestler, Frieda Kage, Georgi Dimchev, Anika Steffen, Theresia E.B. Stradal, Franz Vauti, Hans-Henning Arnold, Klemens Rottner

**Author notes:** **Correspondence:** Klemens Rottner, or Hans-Henning Arnold.

## Abstract

The Arp2/3 complex generates branched actin filament networks operating in cell edge protrusion and vesicle trafficking. Here we employ a novel, conditional knockout mouse model permitting tissue- or cell-type specific deletion of the murine *Actr3* gene (encoding Arp3). A functional Actr3 gene appeared essential for fibroblast viability and growth. Thus, we developed cell lines for exploring the consequences of acute, tamoxifen-induced *Actr3* deletion causing near-complete loss of Arp/3 complex function as well as abolished lamellipodia formation and membrane ruffling, as expected. However, Arp3-depleted cells displayed enhanced rather than reduced cell spreading, employing numerous filopodia, and showed little defects in individual cell migration. Reduction of collective cell migration as observed for instance in wound healing assays likely derived from defects in maintaining directionality during migration, while the principal ability to chemotax was only moderately affected. Analyses of actin turnover at the cell periphery revealed reduced actin turnover rates in Arp2/3-deficient cells, clearly deviating from previous sequestration approaches. Most surprisingly, induced removal of Arp2/3 complexes reproducibly increased FMNL formin expression, which correlated with the explosive induction of filopodia formation. Our results thus highlight both direct and indirect effects of acute Arp2/3 complex removal on actin cytoskeleton regulation.

## Introduction

Actin filament (F-actin) assembly and turnover in cells is tightly regulated by various actin monomer and filament binding proteins, with diverse functions. The generation of actin filaments *de novo* is referred to as nucleation, a complex process that received increasing attention over the past two decades (Campellone and Welch, 2010;Dominguez, 2010;Rottner et al., 2017). In living cells, spontaneous nucleation is likely prohibited by kinetic barriers, imposed for instance by actin monomer binding proteins, to avoid de-regulated, ectopic filament assembly. Cells overcome these barriers by different classes of filament assembly factors each operating at specific subcellular locations and time, allowing respective physiologic processes to occur. These F-actin assembly factors include the Arp2/3 complex plus various different activators, formins as well as the Spire/Cobl/Leiomodin type of actin binding proteins thought to enhance nucleation through simple actin monomer clustering (Chesarone and Goode, 2009;Qualmann and Kessels, 2009). Some members of these classes were also found to function cooperatively, as exemplified by Spire- or APC-formin couples combining Spire/APC-mediated nucleation with formin-mediated elongation of generated actin filaments (Kerkhoff, 2011;Breitsprecher et al., 2012), although the precise modes of action of such cooperative complexes are still to be unraveled (Montaville et al., 2014). Other studies have implied antagonistic functions between actin assembly classes, e.g. between Arp2/3 complex and formins (Suarez et al., 2015;Kadzik et al., 2020), with the actin monomer-binding profilin operating as gatekeeper for Arp2/3-dependent versus ‒independent actin assembly (Rotty et al., 2015). This complexity thus requires development of individual and combined loss of function approaches and systematic and careful side-by-side analysis.

The first and best characterized complex considered today as *bona fide* generator of rapidly-growing, barbed ends of actin filaments is the Arp2/3 complex (Goley and Welch, 2006), which amplifies actin filament assembly by branching daughter filaments off mother filaments that both continue to grow. In recent years, the Arp2/3 complex turned out to be of fundamental importance for numerous cellular processes, including actin-based cell edge protrusion during migration and various additional actin assembly processes at both plasma and intracellular membranes (Rotty et al., 2013;Molinie and Gautreau, 2018), as well as specific interactions of viral or bacterial pathogens with their host cells (Welch and Way, 2013). Interestingly, many of these processes, in particular those occurring at the plasma membrane, involve adaptive responses orchestrated by Arp2/3 complex to mechanical forces (Mueller et al., 2017;Akamatsu et al., 2020), as recently emphasized (Papalazarou and Machesky, 2020). Although the canonical complex is comprised of seven subunits throughout eukaryotic cell evolution, expression of functionally diverse or incomplete complexes - even within the same cell type – have also been described (Chorev et al., 2014;Abella et al., 2016). However, our understanding of their physiological relevance is still scarce (Pizarro-Cerda et al., 2017).

Canonical Arp2/3 complexes are activated to form actin filament branches by a family of so called nucleation promoting factors (NPFs), the name giving member of which, WASp, causes the rare, X-linked Wiskott-Aldrich-Syndrome, and contains the WCA-module (**<u>W</u>**H2 plus **<u>C</u>**onnector plus **<u>A</u>**cidic) at its C-terminal end operating in actin binding (W) and Arp2/3 complex activation (CA). Although it comes in various forms, such as multiple copies of W-domains, this module constitutes the minimal sequence necessary for Arp2/3 activation (Campellone and Welch, 2010), and is thus common to all class I NPFs. We generally agree today that the specificity of Arp2/3 complex functions in cells is largely mediated by temporal and spatial control through NPFs, potentially explaining how Arp2/3 complex activity can be precisely coordinated throughout cells and tissues. Activation of Arp2/3 complex in specialized protrusive structures termed lamellipodia, for instance, is generally thought to be mostly accomplished through the Rac GTPase-effector **<u>W</u>**AVE **<u>R</u>**egulatory **<u>C</u>**omplex (WRC) (Krause and Gautreau, 2014;Alekhina et al., 2017;Schaks et al., 2018;Schaks et al., 2019).

Various distinct experimental approaches have previously been utilized to interfere with Arp2/3 complex function directly, including classical approaches like RNA interference (Rogers et al., 2003;Steffen et al., 2006;Nicholson-Dykstra and Higgs, 2008), inhibition by small molecules (Nolen et al., 2009) or even cytosolic Arp2/3 complex sequestration (Machesky and Insall, 1998;Hufner et al., 2001;Koestler et al., 2013). Yet, we are still falling short on tools for complete and reproducible elimination of Arp2/3 complex function in living cells, likely due to its vital role in cells (see below).

Aside from stable, simultaneous RNAi-mediated suppression of Arp2 and ArpC2 (p34) in fibroblasts (Wu et al., 2012), genetic elimination of the ArpC3 (p21) subunit allowed studying its consequences on ES cell-derived fibroblasts, albeit only for a limited number of cell divisions (Suraneni et al., 2012). These studies uncovered both, commonly shared and divergent views concerning Arp2/3 function in lamellipodia formation *versus* migration efficiency or chemotaxis, which could not be resolved in respective follow-up studies (Wu et al., 2013;Suraneni et al., 2015). Here, we introduce the tamoxifen-inducible elimination of Arp2/3 function in several, independently-generated fibroblast cell lines, derived from a mouse mutant harboring a conditional Arp3 allele that can be removed by Cre-recombinase (Papalazarou et al., 2020). As opposed to previous studies employing Rac- or WRC-deficient cell lines (Steffen et al., 2013;Schaks et al., 2018) and in line with recent data analyzing dendritic cell migration in environments of varying complexity (Leithner et al., 2016), our results suggest that efficient, mesenchymal migration does not always involve lamellipodia, but that these structures become particularly relevant during processes requiring plasticity and establishment and/or maintenance of the direction of migration. Moreover, and distinct from previously published work on knockdown or knockout of the Arp2/3 complex (Suraneni et al., 2012;Wu et al., 2012), we confirm here, as already suggested by previous, independent studies (Steffen et al., 2013), that lamellipodia are dispensable for cell spreading. Finally, our analyses uncover that acute elimination of Arp2/3 complex function leads to upregulation of FMNL formin expression, which is largely responsible for the induced burst of filopodia formation.

## Materials and methods

### Mice and cell lines

Mice homozygous for the *loxP*-site flanked *Act3r* allele were generated as described previously (Papalazarou et al., 2020). For generation of fibroblast cell lines used in this study, E13.5-embryos lacking head and intestines were cut into pieces, digested with trypsin and single cells separated by thorough squeezing through a pipet tip before seeding into fibroblast growth medium. Cells were spontaneously immortalized through continuous passaging. Subsequently, cells were transfected with Cre-ERT2-plasmid, kindly provided by Pierre Chambon (Feil et al., 1997), leading to stable expression of the inducible Cre-recombinase variant in the cytoplasm of clonal cell lines selected by puromycin treatment (2μg/ml). Tamoxifen treatment causes nuclear translocation of Cre-recombinase (Metzger and Chambon, 2001), and subsequent disruption of floxed *Act3r* alleles. Three clones with comparable growth rates were selected for further analyses, termed clones 5, 7 and 19 (Lahmann, 2011). Following selection, cell lines were maintained in DMEM, 4.5 g/l glucose (Gibco) supplemented with 10% FCS (Sigma), 2 mM L-glutamine (Gibco), 2 mM penicillin/streptomycin (Gibco), 0.1 mM non-essential amino acids (Gibco), 1 mM sodium pyruvate (Gibco) and 5μg/ml of puromycin (Sigma), unless indicated otherwise, and gene disruption routinely induced by treatment with (Z)-4-Hydroxytamoxifen for 96 hours (stock solution in DMSO/EtOH and used in a final concentration of 2μg/ml).

### Plasmids and transfections

Spontaneously immortalized fibroblast cell lines were transfected with Cre-ERT2 using Lipofectamine 2000 (ThermoFisher Scientific) following manufacturer’s instructions. All other transfections were done with jetPRIME transfection reagent (Polyplus) according to manufacturer’s protocol. For shRNA-mediated knockdown of FMNL2 and FMNL3, corresponding psiRNA-h7SK-GFPzeo vectors (InvivoGen) harboring knockdown sequences 5’-GGAAGTCTGCGGATGAGATAT-3’ (FMNL2) and 5’-GGTGCAGATTCAAGCGTACCT-3’ (FMNL3) were co-transfected in a 1:1 ratio, and subjected to FACS-sorting prior to tamoxifen/vehicle treatment. For FRAP (see below), EGFP-tagged human β-actin (Clontech, Mountain View, CA, USA) was used, as described previously (Steffen et al., 2013), and pRK5-myc-Rac1-L61 obtained from Laura Machesky (Cancer Research UK, Beatson Institute, UK). EGFP-tagged Arp3 was as described (Stradal et al., 2001), and for generation of EGFP-C2-Arp3B plasmid, the mRNA transcript variant 1 encoding murine actin-related protein 3B (*Actr3b*) (Jay et al., 2000) was synthesized (GenScript Biotech) and cloned into pEGFP-C2 vector (Clontech).

### Western Blotting and intensity quantifications

Detergent-soluble cell extracts were routinely used for all Western blots, and prepared as described (Kage et al., 2017). Soluble protein concentrations were measured using Pierce™ BCA Protein Assay Kit (Thermo Scientific) and determined using a MRX microplate reader (Dynatech Laboratories). Western Blotting was performed according to standard procedures. Glyceraldehyde-3-Phosphate Dehydrogenase (GAPDH) levels were used as loading controls. Primary antibodies were: GAPDH (clone 6C5, #CB1001, Calbiochem), p16A (ArpC5A, clone 323H3; homemade, undiluted hybridoma supernatant) (Olazabal et al., 2002), Arp2 (clone FMS69, Sigma; at a concentration of 1 μg/ml), Arp3 (clone FMS338, Sigma, at a concentration of 1 μg/ml), p34 (Arpc2, #ab205718, Abcam, 1:1000 dilution), FMNL2/3 (#ab57963, Abcam, 1:1000 dilution), Rac1/3 (clone 23A8, Upstate, 1:2000 dilution), ß-actin (AC-15, #A1978, Sigma, 1:10000 dilution), Sra-1 (rabbit polyclonal 4955B, 1:2000 dilution) and Nap1 (rabbit polyclonal 4953B, 1:5000 dilution) (Steffen et al., 2004). Polyclonal rabbit antisera to Abi-1 (1:2000 dilution) and ArpC1 (p40, 1:500 dilution) were raised (by Eurogentec Deutschland GmbH, Köln, Germany) against the synthetic peptides PPVDYEDEEAAVVQYNDPYADGDPAWAPKNYI derived from the human Abi-1 sequence and TARERFQNLDKKASSEGGTAAG derived from the human ArpC1B sequence, respectively, and affinity-purified using corresponding peptides immobilized on CNBr‐sepharose 4B (Amersham Biosciences, Sweden). Specificity of the antisera was confirmed by Western blot detection of the endogenous and ectopically expressed GFP‐tagged proteins. As secondary antibodies, we used peroxidase-coupled anti-mouse IgG (#111-035-062, 1:10000 dilution) and anti-rabbit IgG (#111-035-045, 1: 10000 dilution), which were purchased from Dianova. Protein bands were visualized using chemiluminescent, peroxidase substrate (Lumi-Light Western Blotting Substrate #12015200001, Roche) and an ECL Chemocam Imager (Intas). Quantifications of protein levels from exposed Western blot membranes were done as described (Kage et al., 2017).

### Immunolabelings

For immunolabellings, MEFs were seeded subconfluently o/n (unless indicated otherwise) onto coverslips acid-washed and coated with human fibronectin (25 μg/ml, #11051407001, Roche), as described (Dimchev and Rottner, 2018). For spreading assays, fibroblasts were fixed after different times indicated in Fig. 4. For timepoint 0, coverslips were coated with poly-L-lysin (PLL, Sigma, 0.1 mg/ml) followed by three PBS-washes, and fibroblasts centrifuged onto coverslips for 2 minutes at 1200 rpm prior to fixation.

After brief washing with PBS, cells were routinely fixed with 4% paraformaldehyde (PFA) in PBS for 20 min, and permeabilized with 0.1% Triton X-100 (Sigma) in PBS for 60 sec, unless indicated otherwise. For phalloidin stainings, glutaraldehyde (EM Grade) was added to fixative. For vinculin stainings, fibroblasts were permeabilized with 0.3% Triton X-100/PBS for one minute prior to fixation with 4% PFA/PBS for 20 minutes. All solutions were pre-warmed to 37°C.

Focal adhesions were stained with monoclonal anti-vinculin antibody (#V9131, Sigma, 1:250 dilution), centrosomes with polyclonal anti-gamma tubulin antibody (PA1-28042, ThermoFisher, 1:1500 dilution), and WRC at the cell periphery (Fig. S3) using monoclonal anti-Abi-1 (supernatant of clone 8.3, 1:50 dilution, kindly provided by Giorgio Scita, IFOM Milano). Secondary reagents were Alexa Fluor 594-coupled anti-mouse IgG (#A11032, Invitrogen, 1:200 dilution) and Alexa Fluor 488-coupled anti-rabbit IgG (#A11034, Invitrogen, 1:400 dilution). Phalloidins were either ATTO488-coupled (#AD488-81, ATTO-TEC, 1:200 dilution), ATTO594-coupled (#AD594-81, ATTO-TEC, 1:200 dilution) or ATTO390-coupled (#AD390-81, ATTO-TEC, 1:200 dilution), dependent on experiment and the combinations with antibodies. Nuclei were routinely stained with DAPI, already present in the mounting medium (ProLong Gold, #P36941, Invitrogen) or, for SIM-experiments, with TO-PRO-3 (#T3605, ThermoFisher, 1:1000 dilution).

### Conventional fluorescence and superresolution microscopy and FRAP

Images of immunolabelings were acquired on an inverted Axiovert 100TV epifluorescence microscope (Zeiss) using a 40x/1.3 NA Plan-Neofluar oil immersion objective. The microscope was equipped with an HXP 120 light source for widefield fluorescence illumination, and a Coolsnap-HQ2 camera (Photometrics) as well as electronic shutters (Uniblitz Corporate) driven by VisiView software (Visitron Systems GmbH, Puchheim, Germany).

Structured illumination microscopy (SIM) data were captured using a CFI Apochromat TIRF 100x/1.49 NA oil immersion objective (Nikon) on a Nikon SIM-E superresolution microscope equipped with a LU-N3-SIM 488/561/640 laser unit mounted on a Nikon Ti eclipse. Image acquisition was enabled using a NIS-Elements software (Nikon)-driven N-SIM motorized quad band filter combined with N-SIM 488 and 561 bandpass emission filters and a Hamamatsu Orca flash 4.0 LT camera.

Fluorescence recovery after photobleaching (FRAP) experiments were performed as described (Dimchev and Rottner, 2018). Specifically here, vehicle control or tamoxifen-treated MEFs were transfected with EGFP-ß-actin followed by o/n seeding onto fibronectin-coated coverslips, and subjected to FRAP experiments the following day.

### Random migration, chemotaxis and wound healing assays

For random migration assays, DMSO/EtOH- and respective tamoxifen-treated fibroblast clones were seeded subconfluently into μ-Slide 4 Well Ph+ Glass Bottom chambers (Ibidi) coated with human fibronectin (25μg/ml) six hours prior to time-lapse, phase-contrast microscopy.

Prior to chemotaxis assays, respective cell populations were starved overnight in DMEM. Chemotaxis experiments towards 2.5% FCS and 100 ng/ml HGF (#H9661, Sigma) in DMEM were performed with μ-slide chemotaxis chambers (Ibidi) according to manufacturer’s instructions.

Wound healing assays were performed on cell monolayers grown on fibronectin-coated glass bottom dishes upon removal of standardized, wound-creating silicone culture-inserts (Ibidi) or wounding with a disposable pipet tip.

Time-lapse microscopy data were generated on an inverted Axio Observer (Zeiss) utilizing a 10x/0.3NA Plan-Neofluar air objective for random migration and chemotaxis or a 20x/0.4NA LD Achroplan objective for wound-healing assays. Cells were kept at 37°C and a CO_2_ concentration of 7.5 using an incubation chamber (Incubator XL multi S1, Zeiss) connected to a heating and CO_2_ control unit (PeCon), and maintained in full medium (except for chemotaxis assays). Random migration and chemotaxis were acquired over a time-period of 20 hours and at a frame rate of 4/hour, whereas wound-healing data were also recorded for up to 20 hours with a frame rate of 12/hour.

### Data analysis and processing

Numbers of cells with lamellipodia and numbers of filopodia or concave edges per cell were counted manually from phalloidin images. DAPI and gamma tubulin stainings were employed to manually quantify the number of nuclei and centrosomes, respectively. The length of concave edges was also assessed manually but aided by ImageJ Software. Analysis of spreading behavior and quantification of adhesion numbers and sizes were performed as described in (Steffen et al., 2013) and (Kage et al., 2017), respectively.

ImageJ plugins ‘manual tracking’ and ‘chemotaxis tool’ were used to analyze random migration and chemotaxis. DiPer software (Gorelik and Gautreau, 2014) was utilized to investigate direction autocorrelation in random migration experiments (Fig. 6B).

Collective migration speed of control and tamoxifen-treated fibroblast populations in wound healing assays was assessed as follows: In order to obtain the distance migrated by the cell monolayer, the area engaged was measured at the beginning and after 720 minutes (12 hours) of the wound closure process, as described previously (Steffen et al., 2013), and divided by the width of the camera chip to normalize for employment of distinct setups in the course of time. Migration distances from individual experiments were then divided by time to obtain migration velocities, averaged and displayed as bar charts.

Data obtained from FRAP experiments were analyzed as follows: Half-times of recovery were derived from fluorescence recovery curves as previously described (Koestler et al., 2013;Steffen et al., 2013;Dimchev and Rottner, 2018). Due to the comparably narrow lamellipodia formed by these fibroblasts and in control conditions, fluorescence intensities within a region of 1 μm at the lamellipodium front of control or the corresponding cell periphery of tamoxifen-treated cells were measured using MetaMorph Software (Molecular Devices). Fitted curves were generated in Sigma plot 12.0 (Systat Software) by applying dynamic curve fits for exponential rise to maximum using *f*(*x*) = *y*_0_ + *a* *(1 − *e*^−*b*x*^).

For calculation of the treadmilling factors (TMF) (Lai et al., 2008), the 1 μm region of intensity measurements described above was subdivided into two regions of identical depths (0.5 μm each), referred to as front and back (see Figs 3 and S6). Two separate recovery curves for front and back regions were generated and, in case of control cells, fitted employing a dynamic curve fit for exponential rise to maximum: *f*(*x*) = *y*_0_ + *a* *(1 − *e*^−*b*x*^) for data derived from the front part, and following a sigmoidal curve: 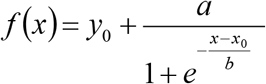 for data derived from the rear part. In contrast, recovery curves from tamoxifen-treated cells were best fitted using equation *f*(*x*) = *y*_0_ + *a* *(1 − *e*^−*b*x*^) for both front and back region. The treadmilling factor was derived from these curves as described (Lai et al., 2008). MetaMorph Software was used to adjust brightness and contrast levels of all images assembled into final figure panels employing Adobe Photoshop CS5 software. Raw data were analyzed and processed in ImageJ, MetaMorph, Excel 2016 (Microsoft) and Sigma plot 12.0.

### Statistics

Statistical significance was assessed using t-test or non-parametric Mann-Whitney rank sum test (Sigma plot 12.0) if datasets were not normally distributed (according to Anderson-Darling normality test), as indicated in individual figure legends. Statistical significance is expressed by the number of asterisks, with * corresponding to a p value smaller than 0.05, and ** and *** corresponding to p<0.01 and p<0.001, respectively.

## Results and discussion

### Acute removal of Arp3 suppresses the Arp2/3 complex and lamellipodia formation

In our initial efforts to generate fibroblast cell lines completely devoid of the Arp2/3 complex subunit Arp3, MEFs homozygous for the conditional *Actr3* allele described previously (Lahmann, 2011;Papalazarou et al., 2020) were both SV40-LT-antigen- or spontaneously immortalized, and subjected to clone isolation and analysis following expression of Cre-recombinase. However, in spite of numerous attempts to isolate homozygously-deleted *Actr3* clones, as successfully achieved for instance in case of the Rho-GTPase Rac1 (Steffen et al., 2013), we were unable to obtain viable *Actr3* null MEFs (data not shown and see below). This indicated that *Actr3* is essential for cell viability and growth, as suggested previously for ArpC2 and Arp2 in fibroblasts, at least in the presence of tumor suppressors (Ink4a/Arf) (Wu et al., 2012), and unlike the recent description of Arp2 null HL-60 promyeloblasts (Graziano et al., 2019). We thus turned to exploring the effects of acute *Actr3* disruption, which was achieved in spontaneously-immortalized cell lines stably expressing tamoxifen-inducible Cre recombinase. Three independent cell clones were isolated (termed Arp3.5, 7 and 19) and analysed separately for Arp3 expression after 3, 4 and 5 full days of treatment with tamoxifen or DMSO/EtOH as vehicle control (Fig. 1A). This allowed to determine 4 days (96h) of tamoxifen treatment as optimal compromise between Arp3 elimination and cell viability (condition boxed red in Fig. 1A). The severe reduction of Arp3 expression after 96 hours correlated well with reduced expression of other Arp2/3 complex subunits tested, i.e. Arp2, Arp-C1, Arp-C2 and Arp-C5 in all three clones (Fig. 1B), confirming the notion that Arp2/3 complex subunits are largely dependent on each other.

**Figure 1.**
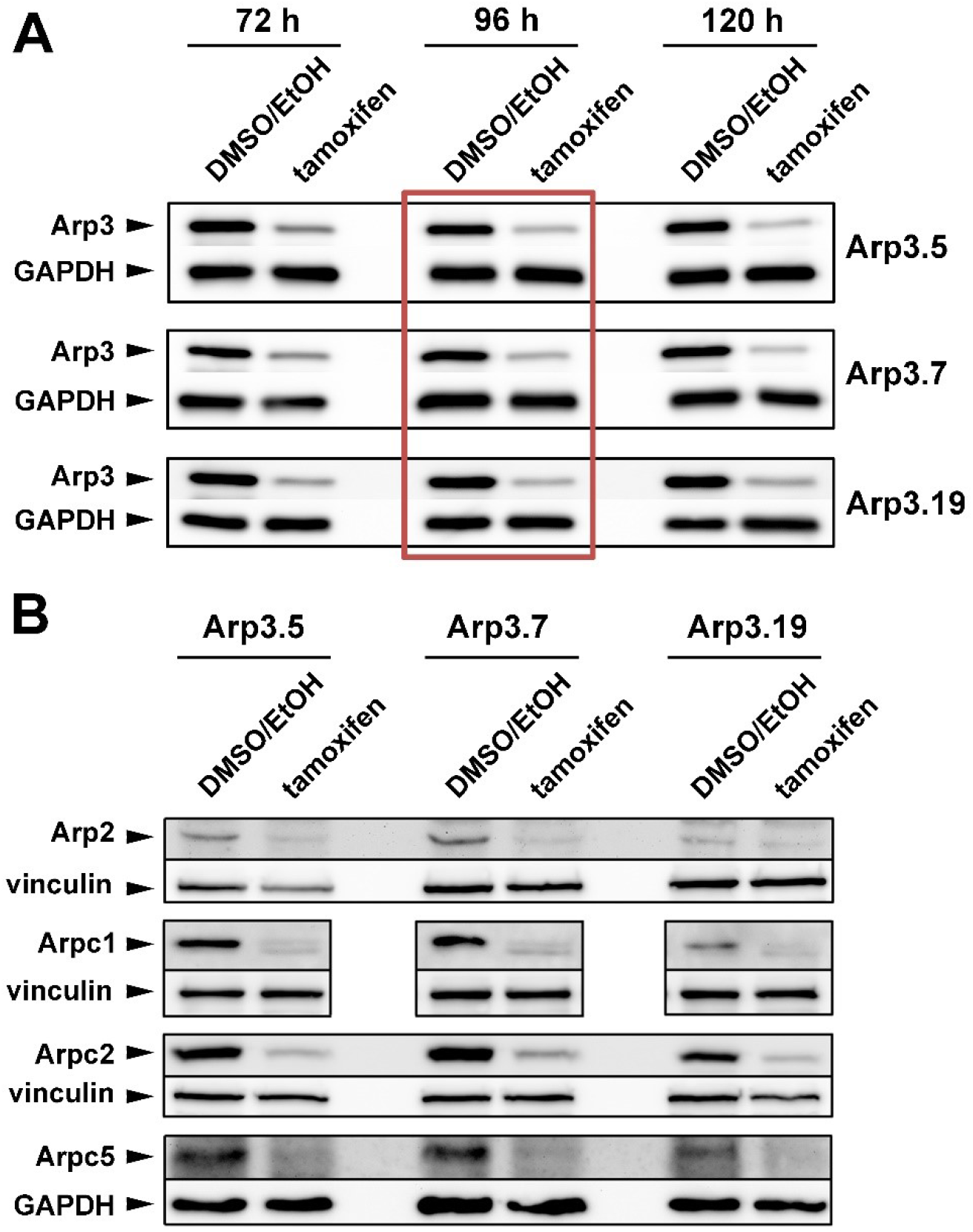
Tamoxifen treatment suppresses expression of Arp2/3 complex subunits. (**A**, **B**) Western blot analysis of Arp2/3 complex subunit levels in individual *Actr3*^*fl/fl*^ clones after tamoxifen treatment (72h, 96h or 120h) or DMSO/EtOH used as vehicle control. GAPDH and vinculin served as loading controls, dependent on the molecular weight of explored Arp2/3 complex subunit. (**A**) Although the extent of Arp3 protein decrease correlated with prolonged tamoxifen treatment, a treatment of 96 hours (red box) was determined as best compromise between efficient protein run down and cell viability, which was significantly compromised upon 120 hours (not shown). The 96h-time point was thus used for all future experiments. (**B**) Tamoxifen-induced removal of Arp3 (96h) concomitantly reduced all additional Arp2/3 complex subunits tested, as indicated.

**Figure 2.**
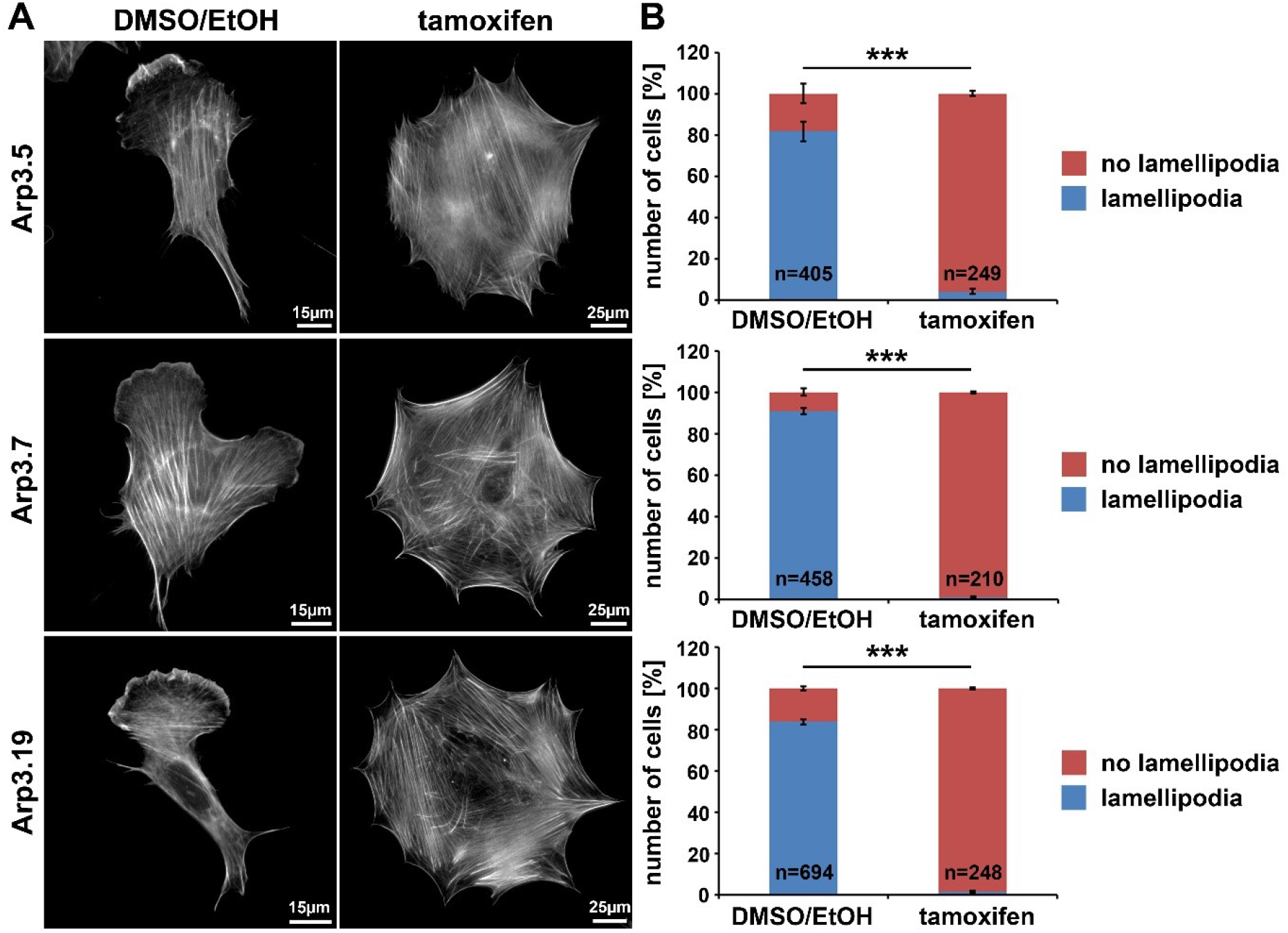
Arp3-depleted MEFs are devoid of lamellipodia. **(A)** Phalloidin-stained examples of distinct *Actr3*^*fl/fl*^ clones (Arp3.5, Arp3.7 and Arp3.19) stably expressing tamoxifen-inducible Cre recombinase, with (tamoxifen) or without (DMSO/EtOH) tamoxifen induction (96h) causing disruption of the *Actr3* gene. Tamoxifen treatment causes near to complete elimination of lamellipodia in representative cells shown. Scale bars as indicated in individual panels. Note the strong increase of spread cell area upon Arp3 disruption. **(B)** Quantification of lamellipodia formation efficiency upon treatments as shown in A. Data are arithmetic means and standard errors of means from three independent experiments; n= total number of cells analyzed; data were statistically compared using two-sided, two-sample t-test (***p<0.001).

**Figure 3.**
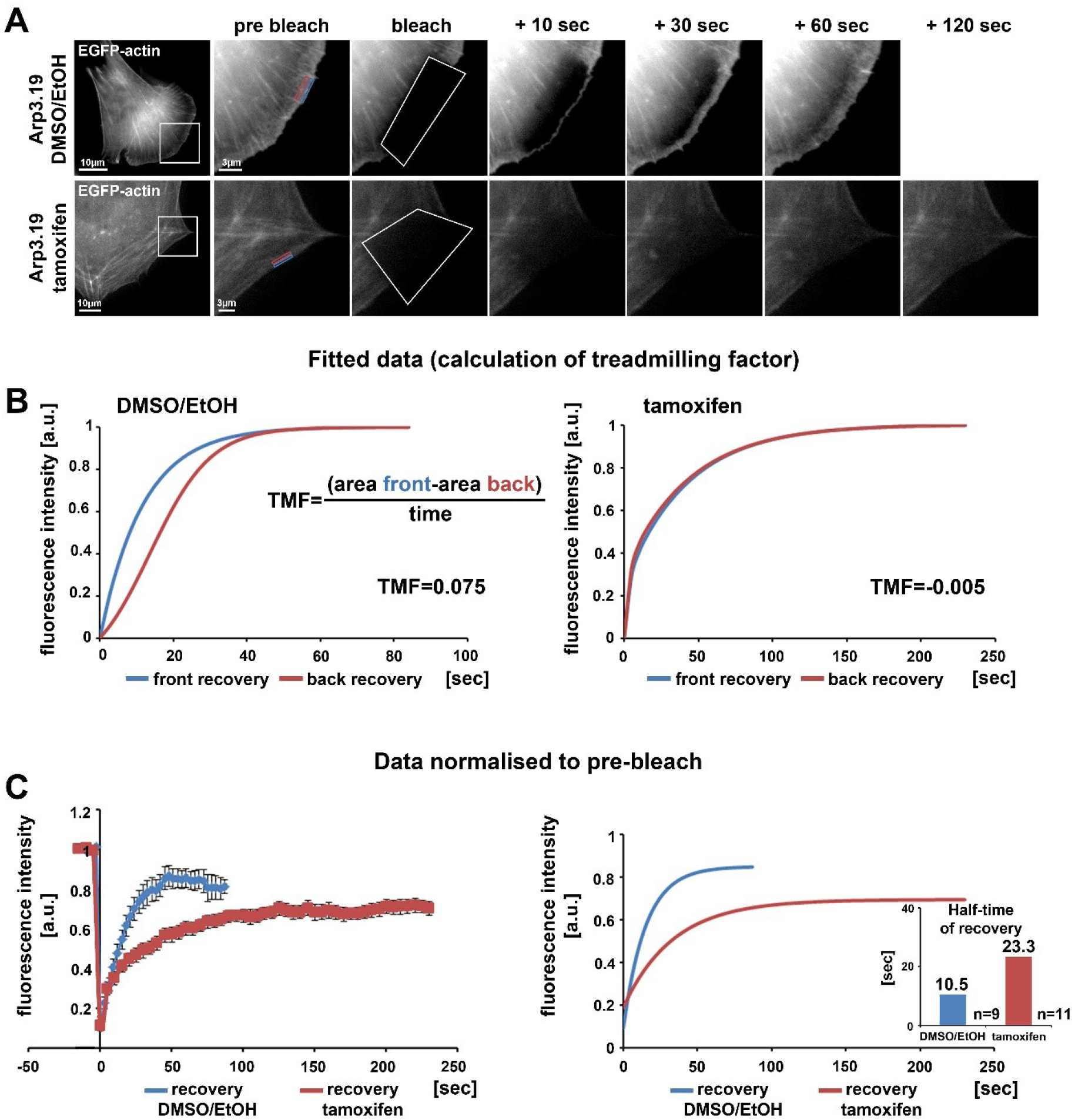
Loss of the Arp2/3 complex interferes with treadmilling and F-actin turnover at the cell periphery. **(A)** Time-lapse images of representative FRAP experiments in *Actr3*^*fl/fl*^ cells with or without tamoxifen treatment (clone Arp3.19) expressing EGFP-actin. Regions of interest (ROI) are marked with white rectangles in overview images on the left. Different time points of enlarged ROI on right-hand panels display fluorescence signals of EGFP-actin immediately pre and after bleach as well as at different time points of fluorescence recovery. White polygons highlight photobleached regions at the cell periphery. Changes in signal intensities over time for calculations of treadmilling factors (TMF) were quantified within a region of 0.5 μm at the front part (blue rectangle) as well as back part (red rectangle) of the lamellipodium in the *Actr3*^*fl/fl*^ cell or the corresponding periphery of the cell following Arp3 depletion. **(B)** Average fitted recovery curves of front (blue) and back (red) regions of Arp3-expressing (left) and Arp3-depleted cells (right) used to calculate the TMF utilizing the illustrated equation. Data were collected from 9 control and 11 tamoxifen-treated cells acquired in three independent experimental days. The treadmilling factor is defined as average difference in fluorescence recovery in the distal *versus* proximal half of the lamellipodium. Note that this difference is absent in tamoxifen-treated cells, demonstrating that presence of a functional Arp2/3 complex is essential for *bona fide* treadmilling at the cell periphery. **(C)** EGFP-actin FRAP recovery curves within a 1 μm deep peripheral region of DMSO/EtOH (blue) *versus* tamoxifen-treated (red) Arp3.19 cells. Analysis was performed on the same time-lapse movies as used for the analysis in (B); data are arithmetic means and SEMs from movies the pre-bleach intensities of which were normalized to 1. Right panel shows fitted curves derived from raw data depicted on the left. Bar chart displays half times of fluorescent recovery as calculated from fitted curves. Note the marked decrease of F-actin turnover (increase of half-time of EGFP-actin fluorescence recovery) upon Arp3 depletion.

**Figure 4.**
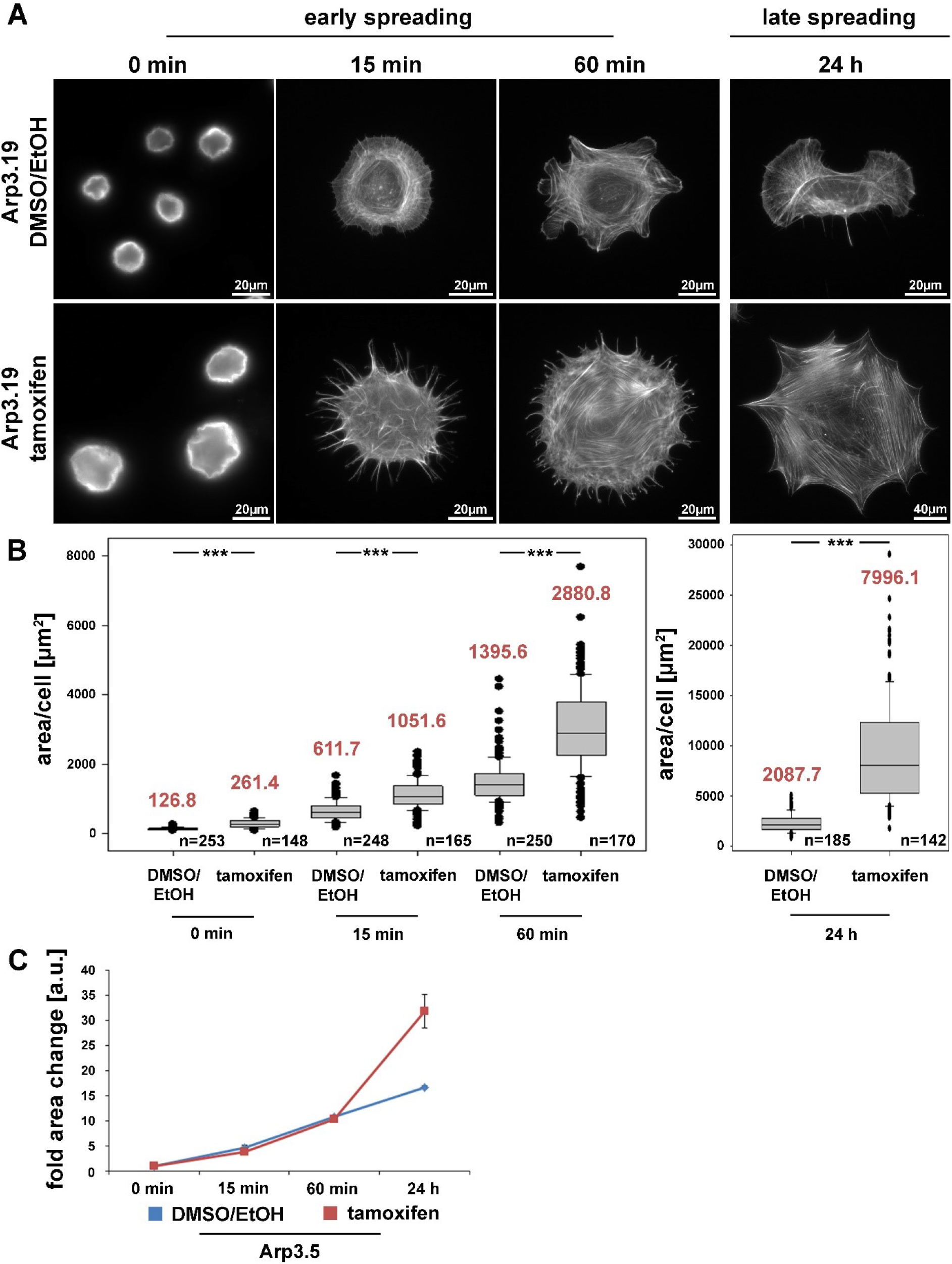
The Arp2/3 complex is not required for cell spreading. **(A)** Phalloidin stainings of *Actr3*^*fl/fl*^ cells (clone Arp3.19) with or without tamoxifen treatment (96h) and subjected to cell spreading for different time points (0 min, 15 min, 60 min and 24 hours). Except for the 0-min time point, for which poly-L-lysine was used (see Methods), cells were allowed to spread on 25 μg/ml fibronectin. **(B)** Box and whiskers plots displaying quantification of spreading area using images as shown in (A). Boxes include 50% (25-75%) and whiskers 80% (10-90%) of all measurements. Outliers are shown as dots. Median values are given in red. n= total number of cells analyzed from three independent experiments. Differences in average cell area of control and tamoxifen-treated cells at different time points of spreading were confirmed to be statistically significant using non-parametric Mann-Whitney rank sum test (*** p<0.001). **(C)** Spreading kinetics of *Actr3*^*fl/fl*^ (EtOH/DMSO, blue) and tamoxifen-treated cells (red) reported as fold change after normalization to cell size at time point 0. Data taken from (B). Data are arithmetic means and error bars represent SEMs. Note that *Arp3* knockout cells spread with kinetics highly similar to corresponding control cells, but adopt a much larger area 24 hours after seeding, due perhaps to continuous increase in cell size effected by Arp3 removal.

To start examining the consequences of acute Arp3 suppression, we quantified the frequency of lamellipodium formation, one of the most prominent structures considered to require functional Arp2/3 complexes (Suraneni et al., 2012;Wu et al., 2012;Suraneni et al., 2015;King et al., 2016). Not unexpectedly, lamellipodia formation was strongly suppressed in all three clonal fibroblast cell lines growing on fibronectin (Fig. 2A, and 2B for quantification). Notably, aside from the elimination of lamellipodia, cells were also significantly increased in size under these conditions (note size bars in Fig. 2A), suggesting that loss of lamellipodia does not necessarily interfere with the expansion of cell area on 2D-surfaces (see also below). Canonical lamellipodia formation is well established to require Rac GTPase signalling to WRC-mediated Arp2/3 complex activation (Steffen et al., 2013;Schaks et al., 2018;Rottner and Schaks, 2019), so to confirm that loss of lamellipodia in these conditions was due to lack of Arp2/3 complex expression, and not due to loss of upstream components of the signalling pathway, several control experiments were performed. 96 hours of tamoxifen treatment did not reduce Rac expression in any of the cell lines employed, as shown using an antibody cross-reactive with Rac1 and −3 (Steffen et al., 2013); instead, a trend towards increased Rac1/3 expression was observed, statistically significant at least in two out of three cell lines (Fig. S1). Interestingly, the Sra-1/PIR121 and Nap1 subunits of the Rac downstream effector WRC were virtually unchanged in expression, whereas the Abl interactor Abi-1 was reduced to about half of controls, for unknown reasons (Fig. S2). However, immunolabeling experiments confirmed the presence of Abi-1 at the cell edges of both vehicle control and tamoxifen-treated cells (Fig. S3), strongly suggesting that the loss of lamellipodia in tamoxifen-treated cells was not caused by the elimination of Abi-1 or other WRC subunits. Finally, the loss of actin filament-rich lamellipodia also did not correlate with a reduction in actin expression, at least as assessed for the prominent, cytoplasmic β-actin isoform (Fig. S4). All this suggested that the loss of lamellipodia upon 96h tamoxifen treatment was caused by suppression of Arp3 and consequently Arp2/3 complex. As final confirmation of this, we found transfection with EGFP-tagged Arp3 to rescue lamellipodia formation in tamoxifen-treated cells in a fashion that was similar in extent to transfection with the related Arp3B (Fig. S5). Notably, using antibodies that could clearly distinguish Arp3 from Arp3B encoded in murine cells by the *Actr3b* gene, we found that Arp3B was undetectable at the protein level with and without *Actr3* deletion (Fig. S5A). This confirmed that Arp3B is not relevant for the cell type and experiments used here, and was consistent with the nearly complete loss of lamellipodia formation upon induced, sole *Actr3* deletion (Figs. 1 and S5B).

### Arp3 depletion abrogates both peripheral actin network treadmilling and F-actin turnover

Lamellipodia are driven by continuous activation and incorporation of Arp2/3 complex into the actin filament network polymerizing at the edge (Lai et al., 2008). This actin network treadmilling behaviour in the lamellipodium was previously disrupted in our lab by the alternative approach of acute inhibition of Arp2/3 complex via its sequestration by microinjection of Scar/WAVE1-WCA (Koestler et al., 2013). In those WCA injection experiments, the network treadmilling behaviour was abolished, but the rate of actin turnover was largely maintained (Koestler et al., 2013), quite distinct from genetic removal of Rac, which strongly reduced F-actin turnover at the cell periphery as well (Steffen et al., 2013). Here we demonstrate a nearly complete elimination of actin network treadmilling at the periphery of cells upon suppression of Arp3 expression (Fig. 3A, B), but also a severe reduction of actin network turnover (Fig. 3C for representative clone 19) to more than double of the fluorescence half time of recovery rate in the KO as compared to control. Essentially the same pattern was observed when cells were transfected with myc-tagged, constitutively active Rac1 (Fig. S6), which entirely failed to stimulate lamellipodia formation in Arp3-deficient cells. Highly comparable results were obtained with clones 5 and 7 both with and without stimulation by constitutively active Rac1 (data not shown). Although the differential response of actin turnover at the cell periphery upon Rac1-KO and Arp2/3 complex sequestration was noted and discussed previously, it remained unclear whether it derived from choice of target protein or experimental approach employed (Steffen et al., 2014). The data shown here clearly establish aforementioned differential response not to derive from choice of protein. Although we cannot exclude that the differences in effects observed upon Arp2/3 complex sequestration (Koestler et al., 2013) *versus* depletion may at least in part reflect the differential timing in development of the phenotype (seconds and minutes in case of sequestration *versus* hours and days in case of induced depletion), the data provided here confirm that sustained interference with Rac signalling and downstream lamellipodial Arp2/3 complex activation abrogates both actin network treadmilling and F-actin turnover.

### Arp2/3 complex is not required for fibroblast cell spreading

Previous studies unanimously concluded that Arp2/3 complex and Arp2/3 complex-dependent lamellipodia are relevant for cell spreading (Suraneni et al., 2012;Wu et al., 2012). However, this view was challenged by our data on Rac-deficient fibroblasts that are also completely devoid of lamellipodia (Steffen et al., 2013), indicating that previous observations derived from specific, Arp2/3 abrogation-dependent roles in spreading rather than a general, lamellipodia-dependent effect. Interestingly, acute suppression of Arp3 expression dramatically increased cell size, but did not reduce spreading efficiency, as assessed from comparing the cell area covering the substratum surface at various time points after seeding (Fig 4A, B). Even if we normalized to the dramatic increase in cell size after 96h of tamoxifen treatment, no substantial relative difference in spreading was recorded at any time point after seeding, neither for clone 19 (Fig. 4C) nor clone 7 (Fig. S7). Only clone 5 slightly lagged behind at early spreading times if considering differences in cell size (Fig. S8C), but this difference was eliminated when data from all three clones were pooled (Fig. S8D). This strongly suggests that clonal variation in this case might explain the observation rather than a genotype-specific effect. We conclude that by and large, the loss of Arp3 and thus Arp2/3 function in actin remodelling and lamellipodia formation can be compensated for by cells in order to explore new space during spreading. As deduced from earlier research, this process likely involved filopodia formation (Wu et al., 2012;Steffen et al., 2013;Suraneni et al., 2015). To confirm this, we performed phase contrast video microscopy of cells immediately after seeding on fibronectin (Video S1). Whereas wildtype cells spread using both, short filopodia and lamellipodia, Arp3-deficient cells initially protruded numerous, prominent and long filopodia, which then served as tracks for the advancement of cytoplasm in between them (Video S1). Together, these data clearly suggest that at least in fibroblasts, cell spreading can be entirely uncoupled from both lamellipodia formation and Arp2/3 complex function.

### Arp2/3 complex promotes directional migration, but is not essential for chemotaxis

Arp2/3 complex is commonly agreed to be crucial for haptotaxis in which cells follow gradients of ECM components or surface-attached chemoattractants (King et al., 2016;Swaney and Li, 2016), but whether or not Arp2/3-depleted cells display a cell-autonomous defect in chemotaxis has remained controversial (Suraneni et al., 2012;Wu et al., 2012;Wu et al., 2013;Suraneni et al., 2015). In classical wound healing experiments, our three cell lines displayed a significant defect in collective wound closure velocity upon acute Arp3 depletion as compared to EtOH-treated controls (Fig. 5 and Video S2). These data were highly reminiscent of previous data obtained with ArpC3-KO, ES cell-derived fibroblastoid cells (Suraneni et al., 2012). However, we were not able to explain this defect by general reduction of migration efficiency, as migration speed was not or only moderately reduced in random migration assays performed with clones 5 and 7/19, respectively (Fig. 6A). Interestingly, assessment of directional migration, which can be done in a fashion unbiased by cell speed (Dang and Gautreau, 2018), was consistently and dramatically impaired upon Arp3 depletion in all three cell lines (Fig. 6B). So Arp2/3 complex activity in our fibroblasts appears more important for maintaining directionality of migration than for rate of motility, consistent with the finding that the Arp2/3 inhibitor Arpin promotes cell turning (Dang et al., 2013;Dang and Gautreau, 2018). This also fits the observation of more compromised performance concerning directional migration in order to close a wound (Fig. 5 and Video S2). To what extent does the observed inability to maintain directionality of migration in random migration assays affect chemotactic performance in our cell lines both with and without Arp2/3 complex? For this, we probed for chemotactic migration towards HGF and 2,5% serum, as used previously for Rac1-deficient fibroblasts (Steffen et al., 2013). In this previous study, we concluded that cells lacking this prominent upstream GTPase, which is considered essential for WRC- and Arp2/3 complex-driven, lamellipodial actin assembly, are incapable of chemotactic migration (Steffen et al., 2013). Thus, it was reasonable to assume that similar results would be obtained with its prominent, direct downstream effector of actin nucleation, the Arp2/3 complex, in spite of clearly conflicting data in this respect with alternative, Arp2/3 complex-depleted fibroblast cell models (Wu et al., 2012;Wu et al., 2013). Importantly, however, the findings described here largely confirm aforementioned data published by the Bear-lab, as the capability to chemotax was still observed upon Arp3 depletion (Fig. 7, clone 19), with the forward migration index (FMI) being only moderately reduced and the defect being more severe if considering overall chemotactic migration rate (Fig. 7D). Nevertheless, our data also shed more light on this long-standing controversy on the precise function of Arp2/3 complex in chemotaxis, as we do find a partial contribution of the presence of Arp2/3 complex in such assays, even though it does not appear to be obligatory, as concluded from other previous data (Suraneni et al., 2012;Suraneni et al., 2015). Virtually identical results were obtained in independent experiments using clones 5 (Fig. S9) and 7 (Fig. S10).

**Figure 5.**
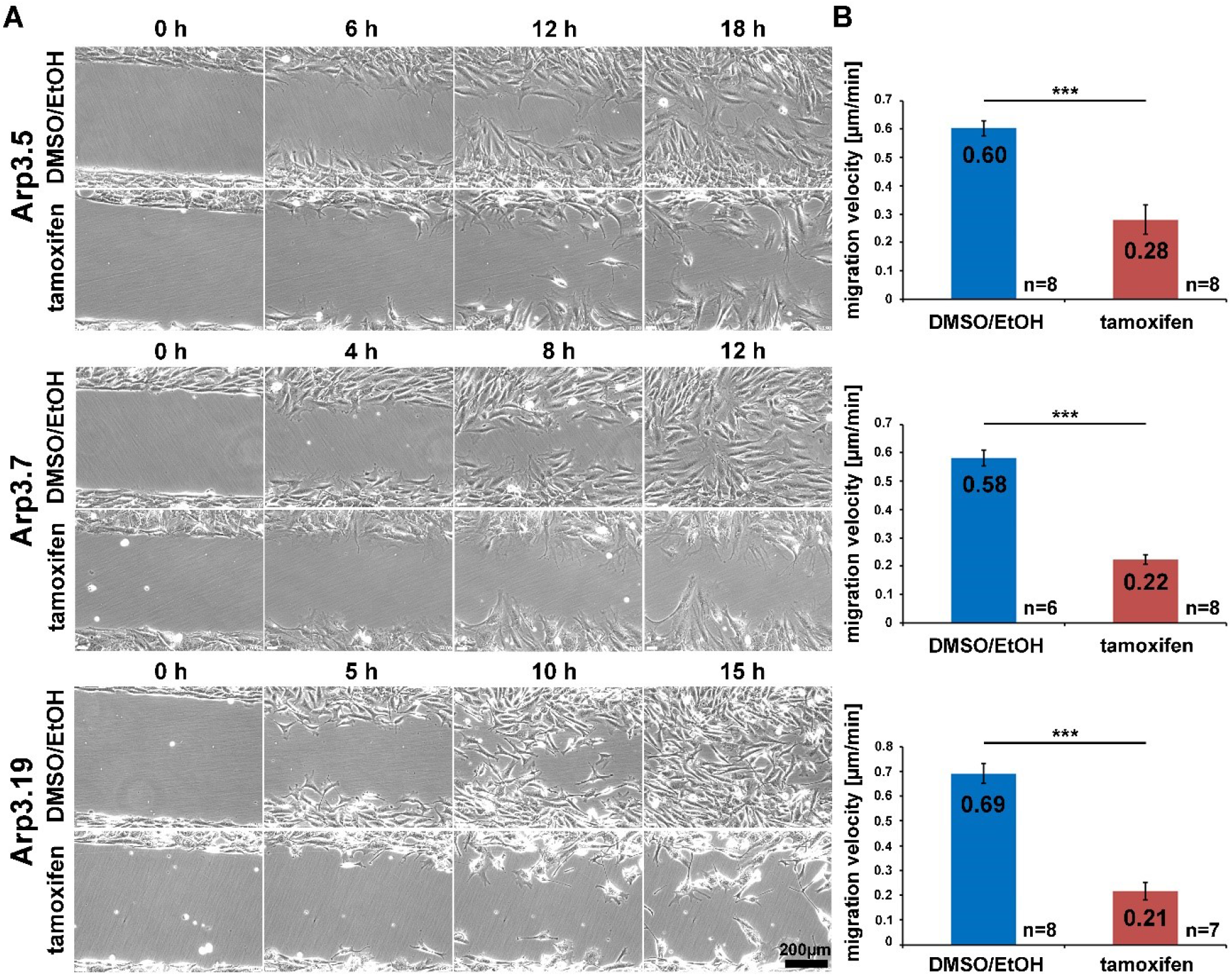
Acute Arp2/3 complex removal reduces wound closure capacity. **(A)** Selected frames from wound healing, time-lapse microscopy (24 hours) of control- and tamoxifen-treated *Actr3*^*fl/fl*^ clones. Wounds closed on average after roughly 18 hours in controls, but failed to do so in Arp3 null cells. **(B)** Quantification of collective migration velocity during wound closure utilizing movies as displayed in (A). Bar charts show arithmetic means and SEMs. n = number of movies analyzed, generated from three independent experiments. Differences in wound closure speed between individual control- and tamoxifen-treated clones were confirmed to be statistically significant using two-sided two-sample t-test (*** p<0.001).

**Figure 6.**
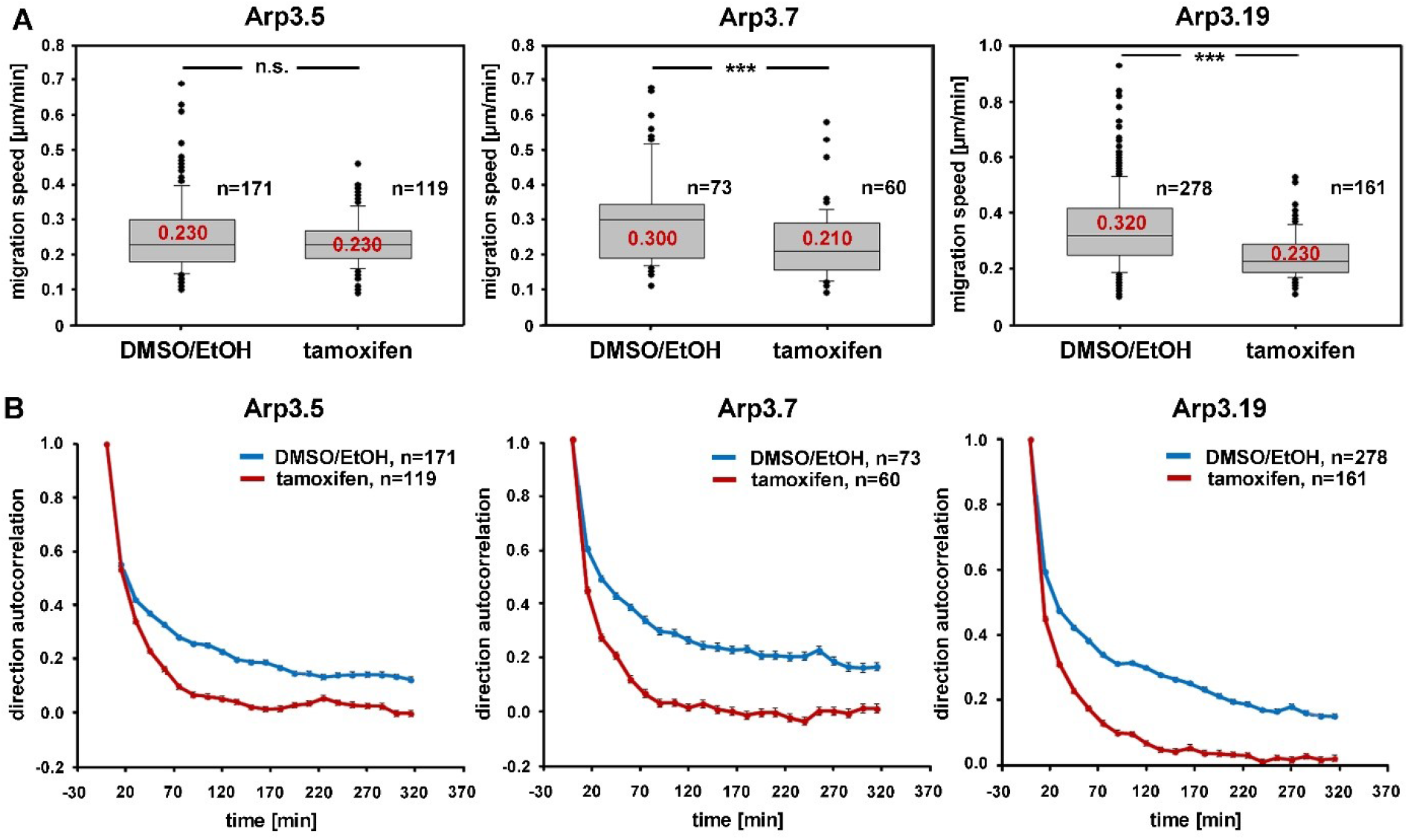
Arp2/3 complex depletion has little effect on the rate of random cell migration, but diminishes its directionality. **(A)** Random migration speed of *Actr3*^*fl/fl*^ clones (Arp3.5, Arp3.7 and Arp3.19, as indicated) with or without treatment with tamoxifen causing acute gene disruption. Data obtained from time-lapse microscopy (20 hours) followed by manual tracking of individual cells. Box and whiskers plots were as described for Fig. 4B, and data derived from three independent experiments. n= total number of cells analyzed. Non-parametric Mann-Whitney rank sum test was used for statistical comparison of respective groups (*** p<0.001). **(B)** Analysis of directional migration utilizing trajectories from (A). An autocorrelation of 1 corresponds to strictly directed movement whereas an autocorrelation of 0 corresponds to pure random walk. Note the consistent loss of directional migration upon Arp2/3 complex removal (tamoxifen) in these experimental conditions in all three clones. Data are displayed as direction autocorrelation curves with error bars representing standard errors of means.

**Figure 7.**
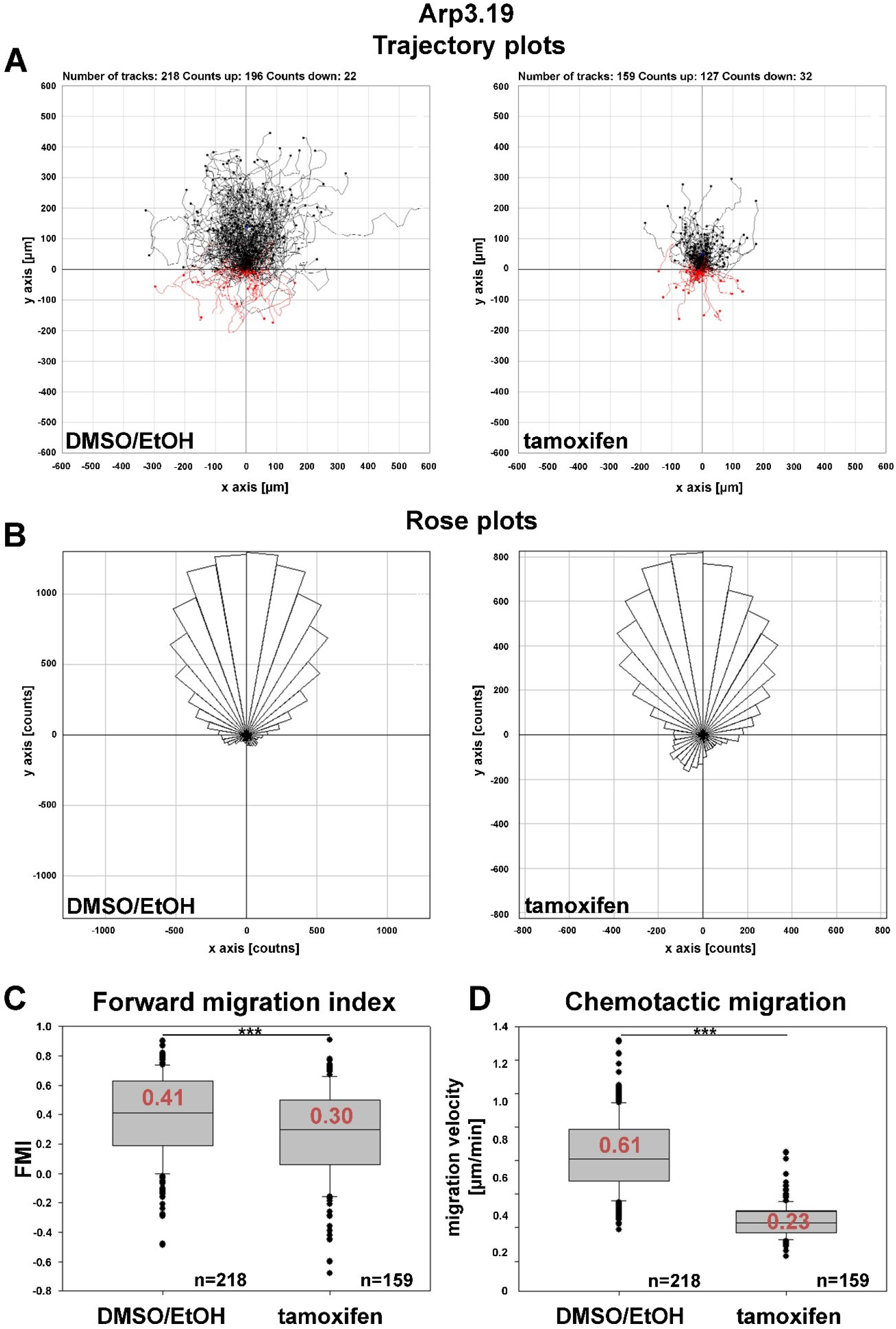
Arp2/3 complex contributes to, but is not essential for chemotaxis. **(A)** Trajectory plots of control (DMSO/EtOH)- and tamoxifen-treated MEFs (*Actr3*^*fl/fl*^ clone Arp3.19) migrating towards a gradient of 2.5% serum and 100ng/ml HGF. Data obtained from time-lapse microscopy (20 hours) followed by manual tracking of cells in three independent experiments. **(B)** Rose plots with each 10° segment showing the frequency of migratory tracks in that particular direction. Note that both DMSO/EtOH- and tamoxifen-treated fibroblasts migrate towards the given gradient with increased frequencies. **(C)** Forward migration index (FMI) obtained from the two experimental conditions. The FMI is defined as displacement of the cell in the direction of the gradient divided by the total distance migrated. Again, Arp3-depleted fibroblasts display a slightly impaired directional persistence. **(D)** Quantification of rates of chemotactic migration. Velocity of chemotactic migration of Arp3-depleted cells is reduced to an extent similar to that observed in directed, wound healing migration assays. Box and whiskers plots in (C, D) were as described for Fig. 4B. Statistics: Non-parametric Mann-Whitney rank sum test (*** p<0.001).

### Defective directional migration coincides with altered focal adhesion patterns

Previous studies have suggested effects of Arp2/3 complex-depletion on certain focal adhesion features, in particular concerning their alignment with each other, which is promoted by functional Arp2/3 complex and the presence of lamellipodia (Wu et al., 2012). Apart from this apparent phenotype (Fig. 8A), we saw a significant increase in overall adhesion numbers per cell (Fig. 8B), but this was reverted to the opposite if the large increase in cell area effected by acute Arp2/3 complex removal was considered (Fig. 8C). Finally, we also found a significant increase in average sizes of individual focal adhesions (Fig. 8D). It is commonly agreed upon that proper focal adhesion turnover is crucial for effective migration. Thus, the strong defects in directional migration that we observed may well be due to a combination of increased cell size and de-alignment as well as increased size of focal adhesions.

**Figure 8:**
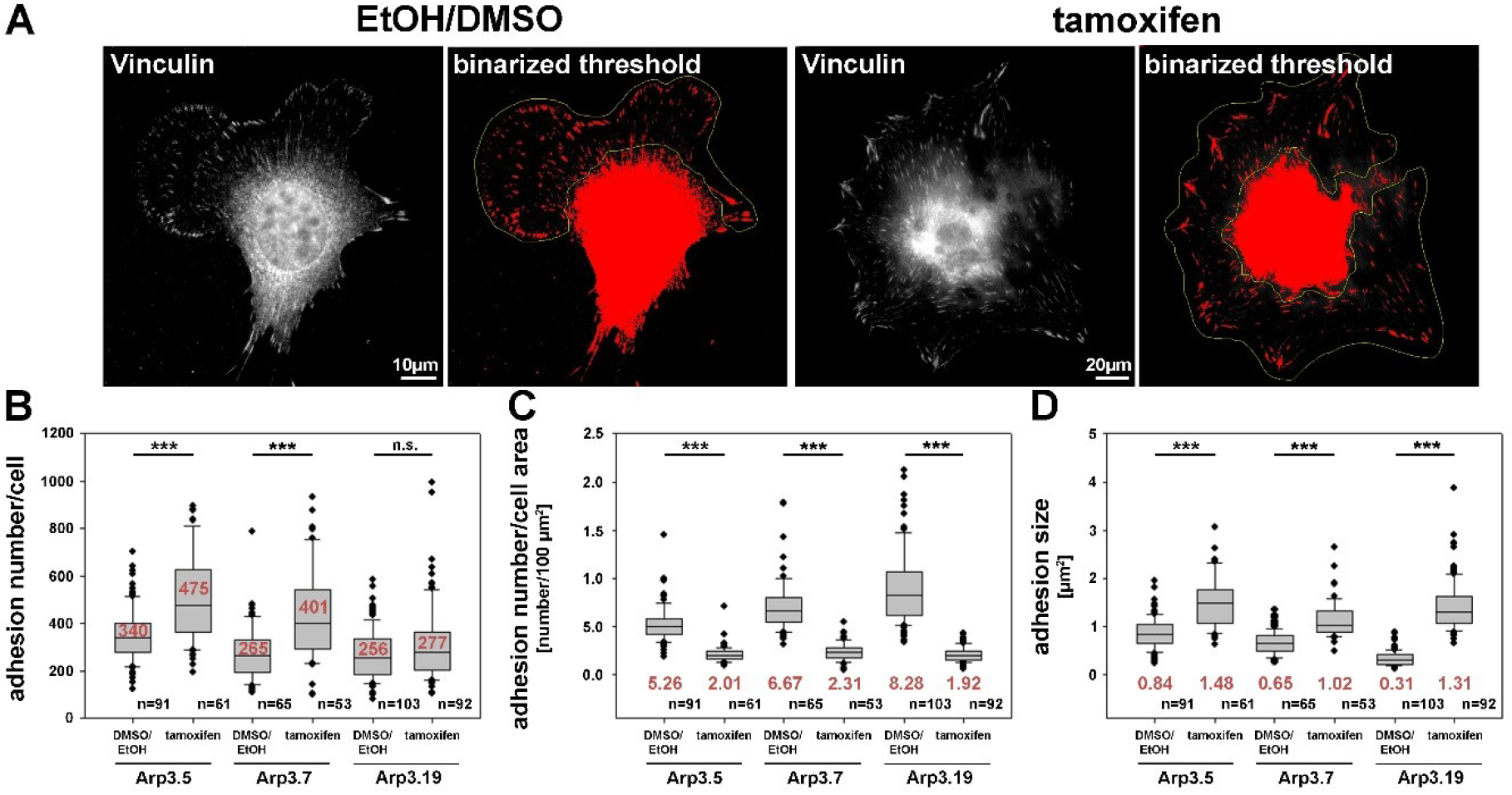
Arp2/3 complex removal alters focal adhesion patterns. **(A)** Representative images of fibroblasts (clone Arp3.19) with or without tamoxifen treatment and stained with anti-vinculin antibodies. A binarized threshold was applied to these stainings in order to facilitate automated analysis of adhesion numbers and sizes within distinct regions of interest (yellow outlines, defined manually). Note the exclusion of the cell centre and perinuclear area due to the lack of staining resolution in these areas. **(B)** Calculation of adhesion number/cell, adhesion number/cell area and **(D)** adhesion size were all based on data obtained from three independent experiments as described in (A). Box and whiskers plots were as described for Fig. 4B. n= total number of cells analyzed. Statistics were performed utilizing the non-parametric Mann-Whitney rank sum test (*** p<0.001). Note that Arp3-deficient cells display more adhesions that are also bigger in size than Arp3-expressing controls. However, the adhesion number per cell area is reduced upon acute Arp2/3 complex removal as compared to control cells.

### Acute Arp3 loss of function increases expression of FMNL subfamily formins that operate in filopodia formation

Our previous studies have established both Arp2/3 complex and FMNL subfamily formins to co-operate in the efficiency of lamellipodia formation (Block et al., 2012;Kage et al., 2017). However, in the absence of lamellipodia, such as upon Rac1 deletion, FMNL formins can only trigger filopodia but not rescue lamellipodia formation (Kage et al., 2017). In these experiments, active FMNL3, which can promote actin filament nucleation independent of Arp2/3 complex *in vitro*, prominently induced the formation of filopodia with club-shaped appearance (Kage et al., 2017), a feature that had previously also been attributed to active actin filament nucleation in cells (Yang et al., 2007;Block et al., 2008). Interestingly, FMNL3 knockdown was recently also found to suppress filopodia formation in U2OS osteocarcoma cells (Young et al., 2018). In a screen for potentially compensatory upregulation of prominent actin binding proteins, both FMNL2 and FMNL3 appeared significantly enhanced in expression upon induced Arp3 suppression (Fig. 9A, B). Since tamoxifen-induced Arp3 depletion clearly increased cell edge complexity, with numerous, small, concave-shaped regions at the cell periphery (Fig. 2A, Fig. S11A), and strongly-increased filopodia numbers (Fig. S11B), we asked whether RNAi-mediated depletion of induced FMNL formins would revert these effects. First, we confirmed the efficiency of combined FMNL2 and −3 knockdown with and without tamoxifen treatment (Fig. 9C). We noted that FMNL formin expression was weak in DMSO/EtOH-treated cells, and knockdown efficiency admittedly moderate. For filopodia quantification, we chose conditions in which filopodia numbers were maximal, which was the case during spreading (Fig. 9D, E). Interestingly, filopodia numbers seen in the presence of Arp2/3 complex in these conditions, were not affected by FMNL2/3 knockdown (for quantification see Fig. 9E), but the increase in filopodia numbers effected by Arp3 removal appeared abolished (Fig. 9E). We should note that additional actin polymerases have previously been implicated in filopodia formation, foremost of all Ena/VASP family members (Dent et al., 2007), although they are certainly not obligatory for these structures (Damiano-Guercio et al., 2020). It is tempting to speculate that during spreading in the presence of Arp2/3 complex, Ena/VASP proteins are more relevant for the formation of the filopodia formed under these conditions than FMNL2/3 proteins. Interestingly, the Ena/VASP members VASP and Mena were also found to be upregulated in expression in a previous Arp2/3-KO model (Rotty et al., 2015), and we were able to confirm this for VASP upon induced Arp3 removal (data not shown). Thus, the precise relative contributions of both FMNL formin and Ena/VASP families to filopodia formation in the presence or absence of Arp2/3 complex remains to be established in the future. Notwithstanding this, we show here that FMNL formin upregulation has clear effects on the changes of features of protrusion and cell morphology seen upon acute Arp3 removal (Fig. 9). In spread cells of regular cultures, filopodia formation was virtually absent in the presence of Arp2/3-complex, but the increase effected by Arp3 removal was again almost entirely eliminated through additional FMNL2/3 knockdown (Fig. S11B). Alongside with this, cell edge complexity was also severely reduced upon FMNL2/3 knockdown in Arp3-depleted cells, as evidenced by a strong reduction of the numbers of concave edges (Fig. S11C), which of course correlated with an increased size of remaining concave edges upon additional FMNL2/3 knockdown (Fig. S11D). Finally, spreading cells (15 minutes) suppressed in expression of both, Arp2/3 complex and FMNL2/3 displayed severe defects in overall cell shape and actin filament distribution at the cell periphery (Fig. 9D bottom right), nicely illustrated by SIM superresolution imaging (Fig. S12 bottom right panel). Both epifluorescence (Fig. S13A) and SIM imaging (Fig. S13B) in Arp3-KO/FMNL2/3 knockdown cells often showed distinct, oval-shaped regions of very low F-actin content and likely mediating cell-substratum adhesion, which did not coincide with the shape of the nucleus (Fig. S14). Most strikingly, 3D-projections of SIM imaging data of control (Video S3) *versus* Arp3/FMNL2/3-depleted cells (for representative examples without and with nucleus see Videos S4/5 and 6, respectively) clearly revealed a drastic shape change upon loss of both Arp3 and FMNL2/3 formins, the latter of which frequently adopted the shape of thick disks during spreading instead of the flat cone-shaped cells observed in control cells. The physics behind this phenomenon requires further, thorough investigation, including potential effects on stability and nature of the actin cortex in these conditions (Bovellan et al., 2014). Notwithstanding this, our data unequivocally show that the drastic changes in cell morphology effected by acute Arp3 removal, e.g. the observed increase of the numbers of filopodia or of concave regions formed along the cell edge, are to a large extent mediated by the induced upregulation of FMNL2/3 formin expression.

**Figure 9:**
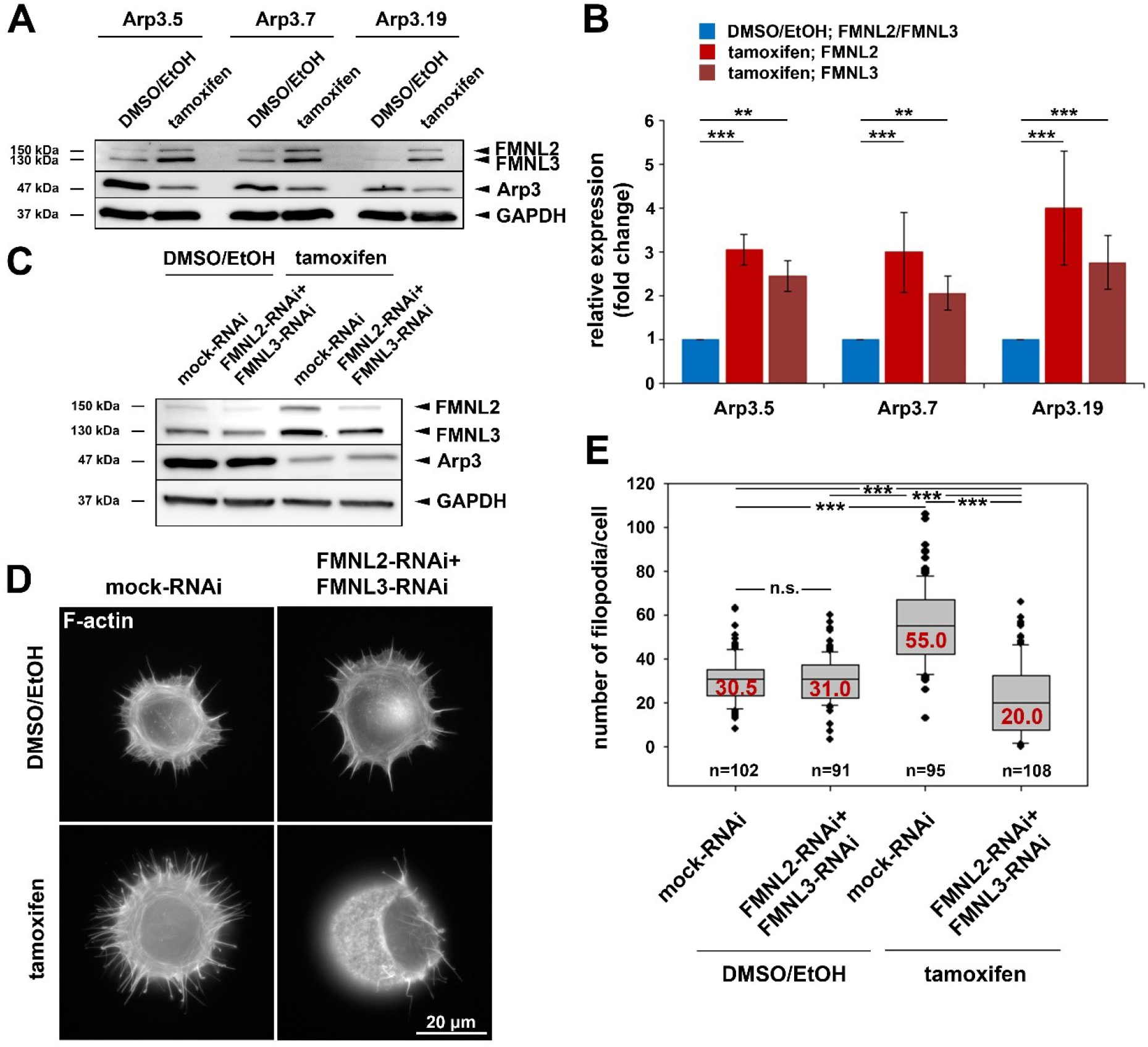
Acute Arp3 removal up-regulates FMNL formin expression enhancing filopodia formation. **(A)** Representative Western blot of Arp3 as well as FMNL2 and FMNL3 protein levels in *Actr3*^*fl/fl*^ clones with and without tamoxifen treatment, as indicated. GAPDH served as loading control. **(B)** Quantification of FMNL2 and FMNL3 protein levels from Western blots as shown in (A). Bar charts display arithmetic means of FMNL2 and −3 protein levels normalized to GAPDH, with each FMNL variant (shades of red) being presented as fold-change to its respective control (blue). Error bars represent SEMs; data obtained from at least five independently generated cell extracts. Statistics were performed utilizing non-parametric Mann-Whitney rank sum test (*** p<0.001; **<0.01). Note the significant elevation of FMNL2 and −3 protein levels upon Arp3 suppression in all three clones. **(C)** Western blot analysis of FMNL2 and FMNL3 protein amounts in DMSO/EtOH- and tamoxifen-treated *Actr3*^*fl/fl*^ cells (clone Arp3.19) after co-transfection with RNAi plasmids individually downregulating FMNL2 and FMNL3 expression or mock-plasmid as control, as indicated. Again, note the elevation of FMNL2 and −3 levels upon Arp3 depletion, which was partially counteracted by combined FMNL2/3 RNAi. **(D)** Representative phalloidin stainings of *Actr3*^*fl/fl*^ cells (clone Arp3.19) control or tamoxifen-treated, and combined with mock- or FMNL2/3-RNAi after 15 minutes of spreading on fibronectin-coated coverslips. **(E)** Quantification of filopodia in cells as presented in (D); box and whiskers plots as described for Fig. 4B. n= total number of cells analyzed from three independent experiments. Statistics were performed utilizing non-parametric, Mann-Whitney rank sum test (*** p<0.001; n.s. not significant). Note that Arp3 knockout is accompanied by a pronounced increase in filopodia numbers that are significantly reduced upon FMNL2/3 RNAi.

### Acute Arp3 removal increases centrosome and nucleus numbers and causes nuclear deformation

As described above, we have been unable to generate clones of immortalized fibroblast cell lines lacking Arp3. This suggested that expression of Arp3 was essential for cell division or perhaps cytokinesis. Indeed, Arp2/3 complex function has previously been linked to proper cell division and mitotic spindle formation (Plessner et al., 2019), perhaps through actin-dependent tuning of centrosomal microtubule nucleation (Farina et al., 2019), although it is already clear that Arp2/3 complex cannot be obligatory for cell division in all cell types and conditions (Graziano et al., 2019). Here we found that induced, acute Arp3 removal in fibroblasts causes dramatic increase in average centrosome numbers from roughly 2 to >5 on average in interphase cells (Fig. 10A-C, Fig. S14, Fig. S15). Severe problems with cytokinesis were also suggested by the increase of average nuclei numbers observed in all three cell clones after acute Arp3 reduction (Fig. 10D). Finally, many nuclei appeared swollen, fragmented or deformed, with frequent “budshape-like” protrusions, which were very rarely seen in control DMSO/EtOH-treated cells. In contrast, the percentage of cells displaying at least one deformed nucleus was increased to roughly 70-80% upon tamoxifen treatment, dependent on the clone treated (Fig. 10B, E). To what extent Arp2/3-dependent actin assembly might contribute to nuclear envelope rupture as previously established in starfish oocytes (Wesolowska et al., 2020) and as pre-requisite for cells to enter metaphase of mitosis, remains to be established in future studies. Clearly, a potential, specific function in nuclear envelope rupture and breakdown would at least be consistent with the deformed nucleus phenotype observed here (Fig. 10B, Fig. S14B, Fig. S15B). However, future research is needed to dissect whether the obligatory function in cell division and growth observed for the *Actr3* gene studied here reflects a cell type- or condition-dependent function in these processes for Arp2/3 complex-dependent actin remodeling or a more general, Arp2/3 complex-independent function of the *Actr3* gene in all cells and tissues (Vauti et al., 2007).

**Figure 10.**
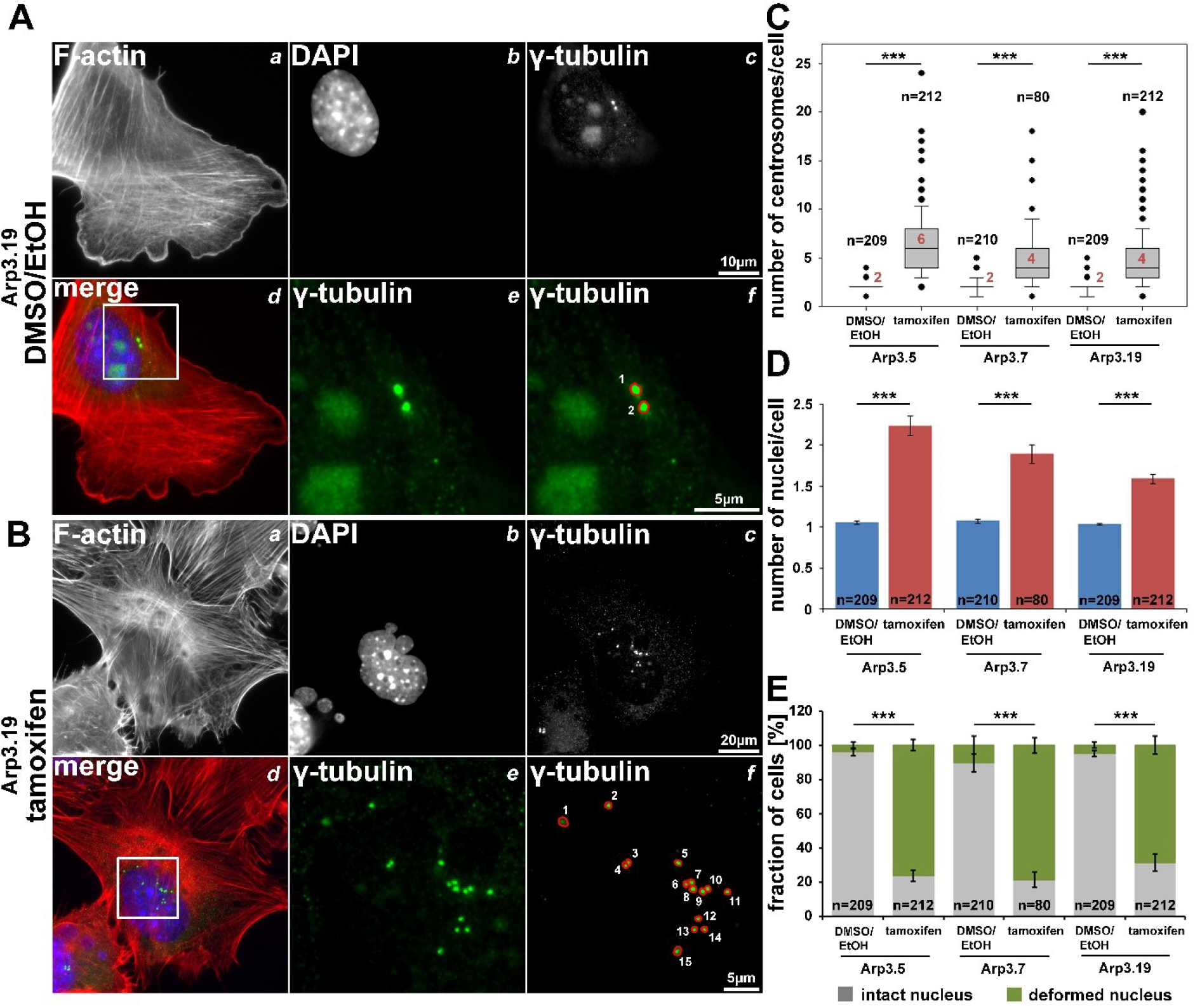
Arp3 depletion causes an increase of centrosome and nucleus numbers as well as nuclear deformation. **(A/B)** Representative fluorescence microscopy images of *Actr3*^*fl/fl*^ cells (clone Arp3.19) stained for the actin cytoskeleton with phalloidin **(a)**, nuclei using DAPI **(b)** and γ-tubulin for centrosomes **(c)**. Merged images **(d)** display actin filaments in red, nuclei in blue and centrosomes in green. White rectangles in (d) mark regions of interest enlarged in **(e)** displaying centrosomes, the quantification of which is illustrated in **(f)**. **(C/D)** Box and whiskers plots (as described for Fig. 4B) displaying numbers of centrosomes (C) or of nuclei (D). For statistics, non-parametric, Mann-Whitney rank sum tests were used (*** p<0.001). **(E)** Analysis of nuclear morphologies. Cells were categorized according to the morphological integrity of their nuclei (intact nuclei in grey; deformed nuclei in green), represented as a fraction of all cells analyzed in individual cell populations. Results are depicted as stacked columns representing arithmetic means and SEMs from three independent experiments; non-parametric, Mann-Whitney rank sum test for statistics (*** p<0.001), n = total number of cells analyzed in (C-E).

## Author Contributions

VD performed all experiments except for the wound healing assays, which were done by SK. FK helped with SIM data generation and cloned FMNL knockdown constructs. IL generated the mouse model harboring loxP site-flanked *Actr3* alleles, established and provided spontaneously immortalized *Actr3*^*fl/fl*^ cell lines and performed initial experiments. GD, AS and TS provided essential reagents and supported data analysis and interpretation. FV aided construct generation for homologous recombination and all experiments involving generation of the mouse mutant. HA conceptualized the Arp3 knockout study, supervised all stages of the project, and helped developing the cellular work. KR supervised all cell experiments, interpreted data and wrote the paper. All authors contributed to data interpretation and development of conclusions, and revised, read and approved the manuscript text.

## Acknowledgments

We thank Sophia Leschik for experimental help during early stages of the project, Pierre Chambon, Laura Machesky and Giorgio Scita for reagents, Lothar Gröbe (HZI Braunschweig) for cell sorting, and Gerd Landsberg (University of Bonn) and Brigitte Denker (HZI Braunschweig) for excellent technical assistance.

## Abbreviations used in this paper

AOTF: Acousto-optical tunable filter
Arp2/3: actin-related protein 2/3
FMNL2: formin-like family member 2
FMNL3: formin-like family member 3
MEFs: mouse embryonic fibroblasts
Rho: Ras homolog gene family
Rac: ras-related C3 botulinum toxin substrate
RNAi: RNA interference
Scar: Suppressor of cAMP receptor
WAVE: WASP family verprolin-homologous
WRC: WAVE regulatory complex
VASP: Vasodilator-stimulated phosphoprotein
Ena/VASP: Enabled/vasodilator-stimulated phosphoprotein
EGFP: Enhanced Green Fluorescent Protein
FRAP: fluorescence recovery after photobleaching

## Supplementary Information

### Legends to supplementary figures

**Figure S1.**
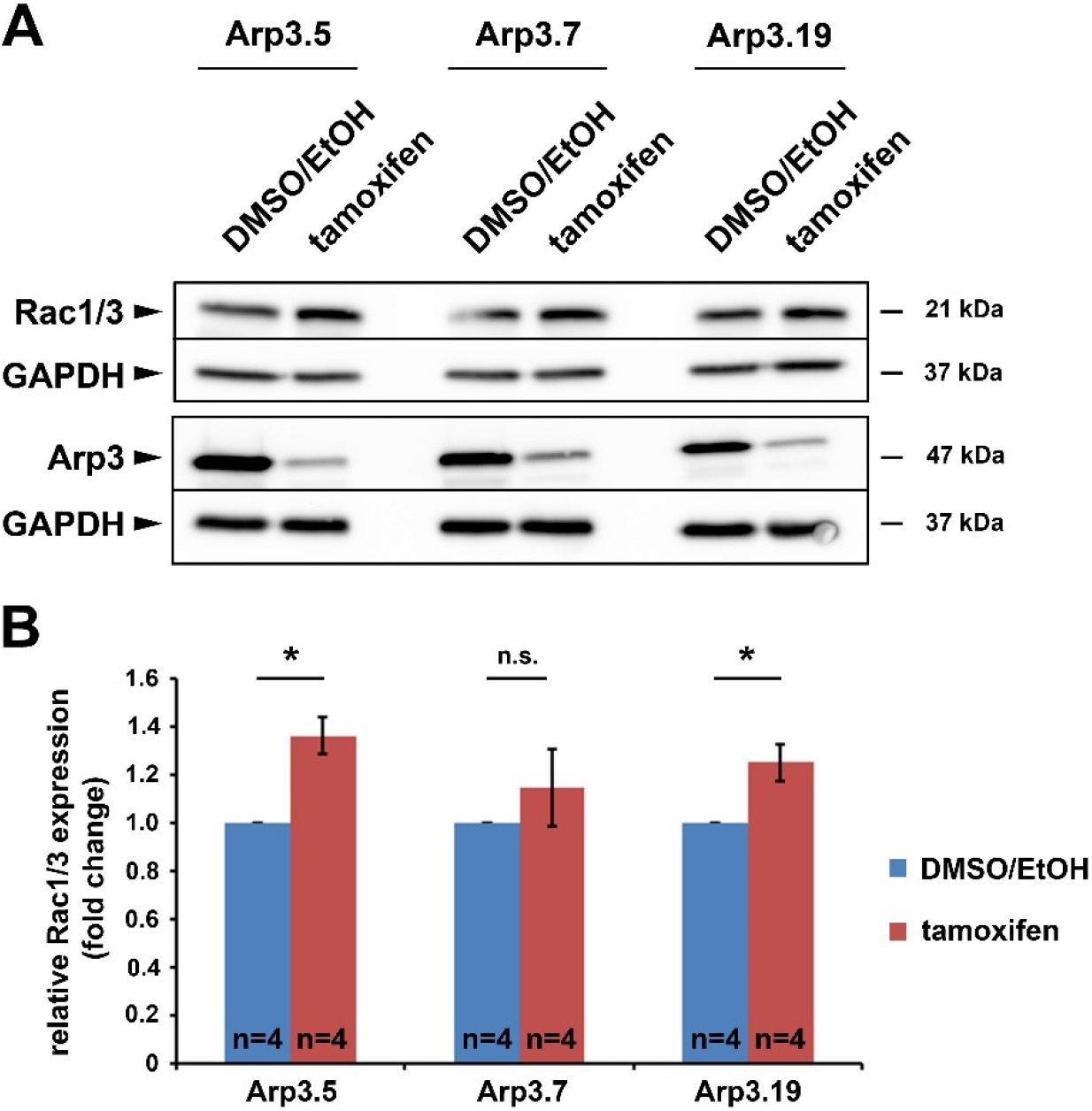
Arp3 depletion increases Rac expression levels. **(A)** Representative Western blot showing Rac protein levels using an antibody cross-reactive with Rac1 and −3 on cell extracts from *Actr3*^*fl/fl*^ cells with and without Tamoxifen treatment. GAPDH served as loading control. **(B)** Quantification of Rac protein levels from Western blots as described in (A). Bar charts display arithmetic means of Rac protein normalized to protein levels of GAPDH, with each knockout cell population (red) being presented as fold-change to its respective control (blue). Error bars represent standard error of means. Data were obtained from four independent experiments. Statistics were performed using non-parametric, Mann-Whitney rank sum test. Increase of Rac levels upon Arp3 removal were confirmed to be statistically significant in Arp3.5 and Arp3.19 cells (* p<0.05), and at least followed the same trend in clone Arp3.7 (n.s. not significant).

**Figure S2.**
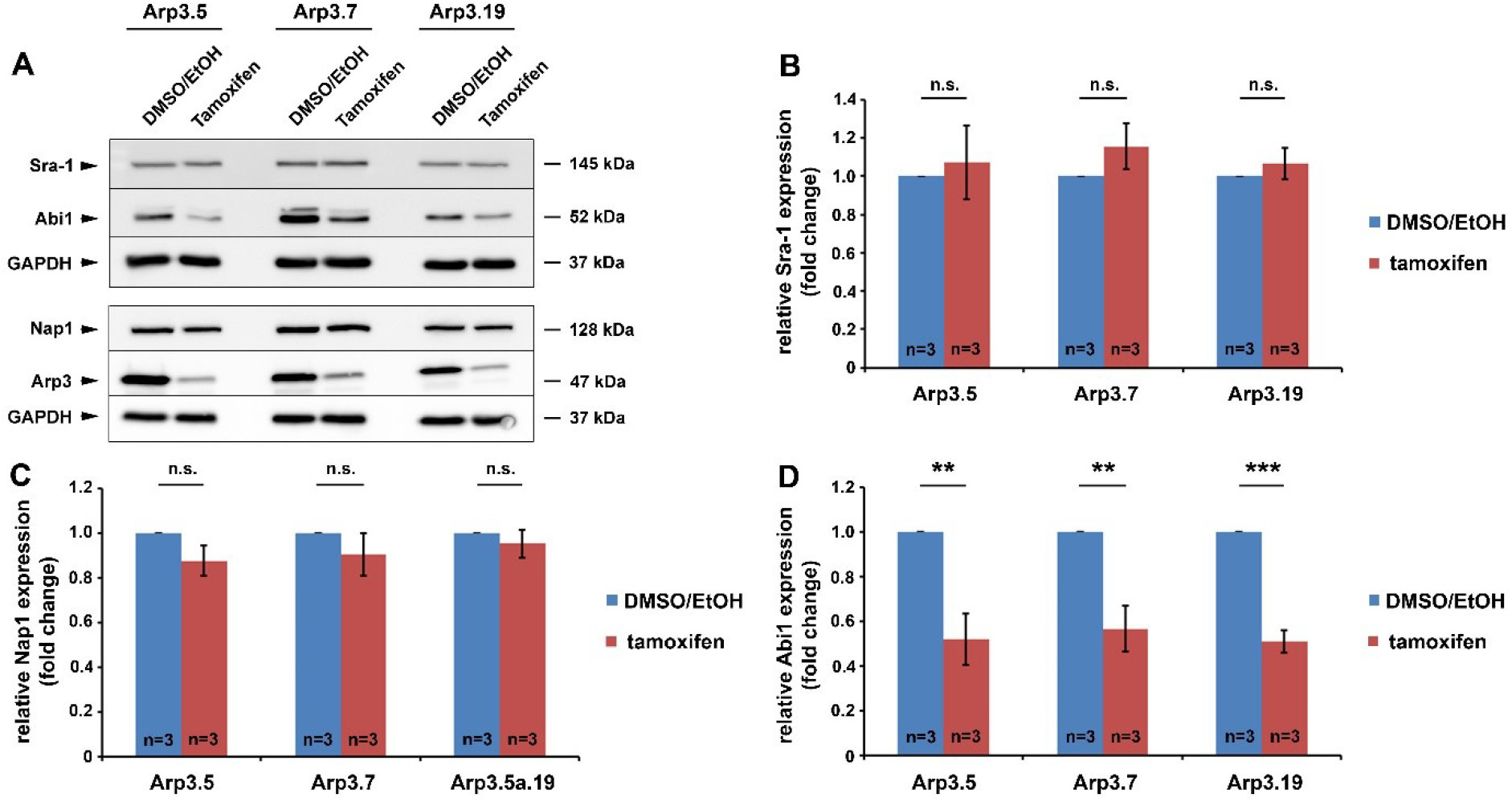
Expression levels of the WRC subunit Abi1 are decreased upon acute Arp3 knockout. **(A)** Representative Western blot for WRC subunits Sra-1, Abi1 and Nap1 in *Actr3*^*fl/fl*^ clones with and without tamoxifen treatment, as indicated. For Nap1 expression, lysates used were identical to the ones used for Rac protein quantification in Fig. S1, hence Arp3 and GAPDH bands are identical. GAPDH served as loading control. **(B/C/D)** Quantification of protein levels from Western blots as shown in (A). Bar charts display arithmetic means +/− SEMs for Sra-1 **(B)**, Nap1 **(C)** and Abi1 **(D)** protein levels, as indicated, and normalized to GAPDH, with each knockout (red) being presented as fold-change to its respective control (blue). Data were obtained from three independent experiments, and statistics performed using non-parametric, Mann-Whitney rank sum test, confirming differences for Abi1 protein levels to be statistically significant (* p<0.05, *** p<0.001, n.s. not significant).

**Figure S3.**
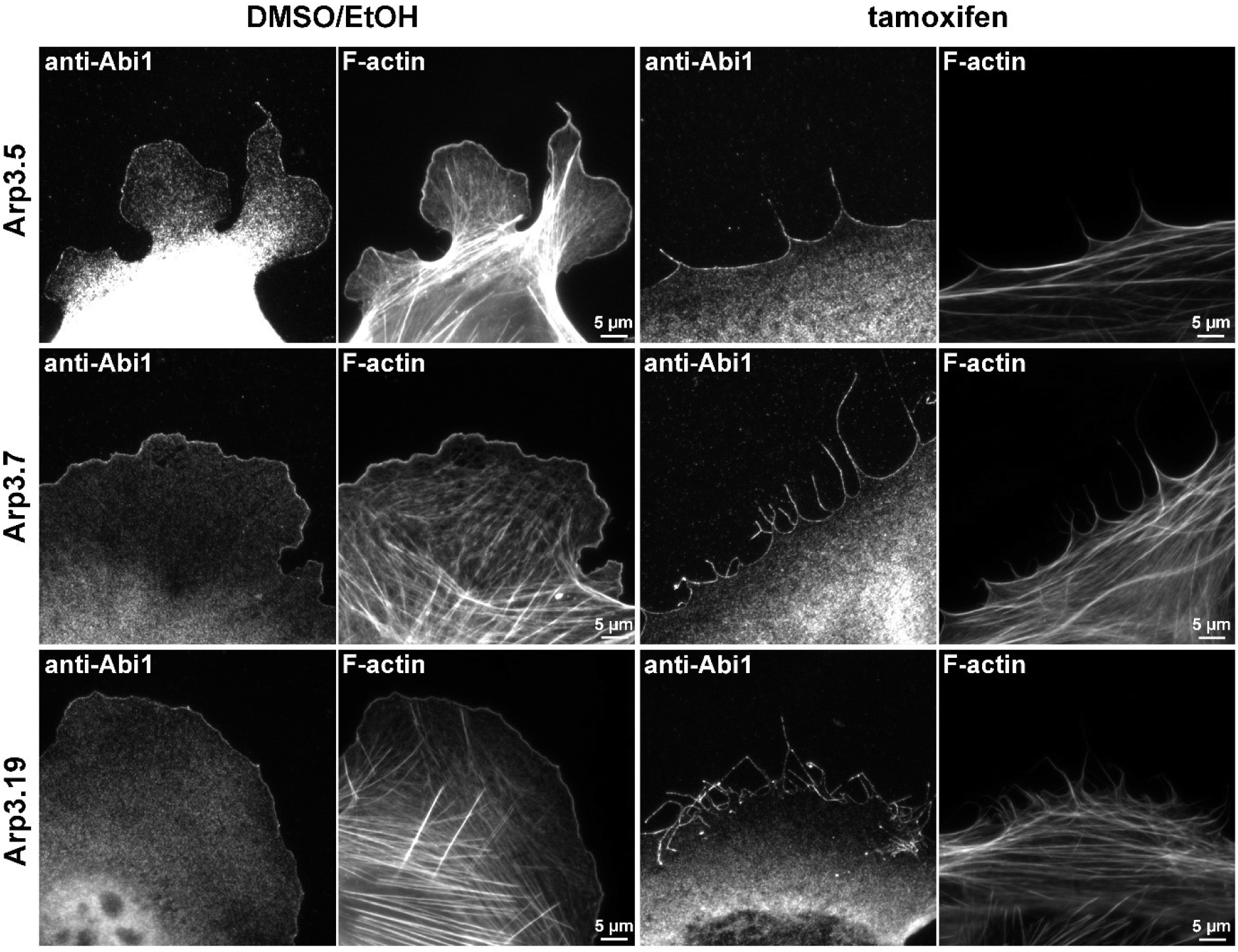
Loss of lamellipodia upon Arp3 removal does not coincide with displacement of Abi from the cell periphery. Immunolabeling with anti-Abi1 antibodies and counterstaining for the actin cytoskeleton of *Actr3*^*fl/fl*^ clones with or without tamoxifen treatment, as indicated. Note the association of the WRC subunit Abi1 with peripheral F-actin bundles despite the absence of lamellipodia in tamoxifen-treated cells, implying that lamellipodia cannot form due to the absence of Arp2/3 complex, and not WRC.

**Figure S4.**
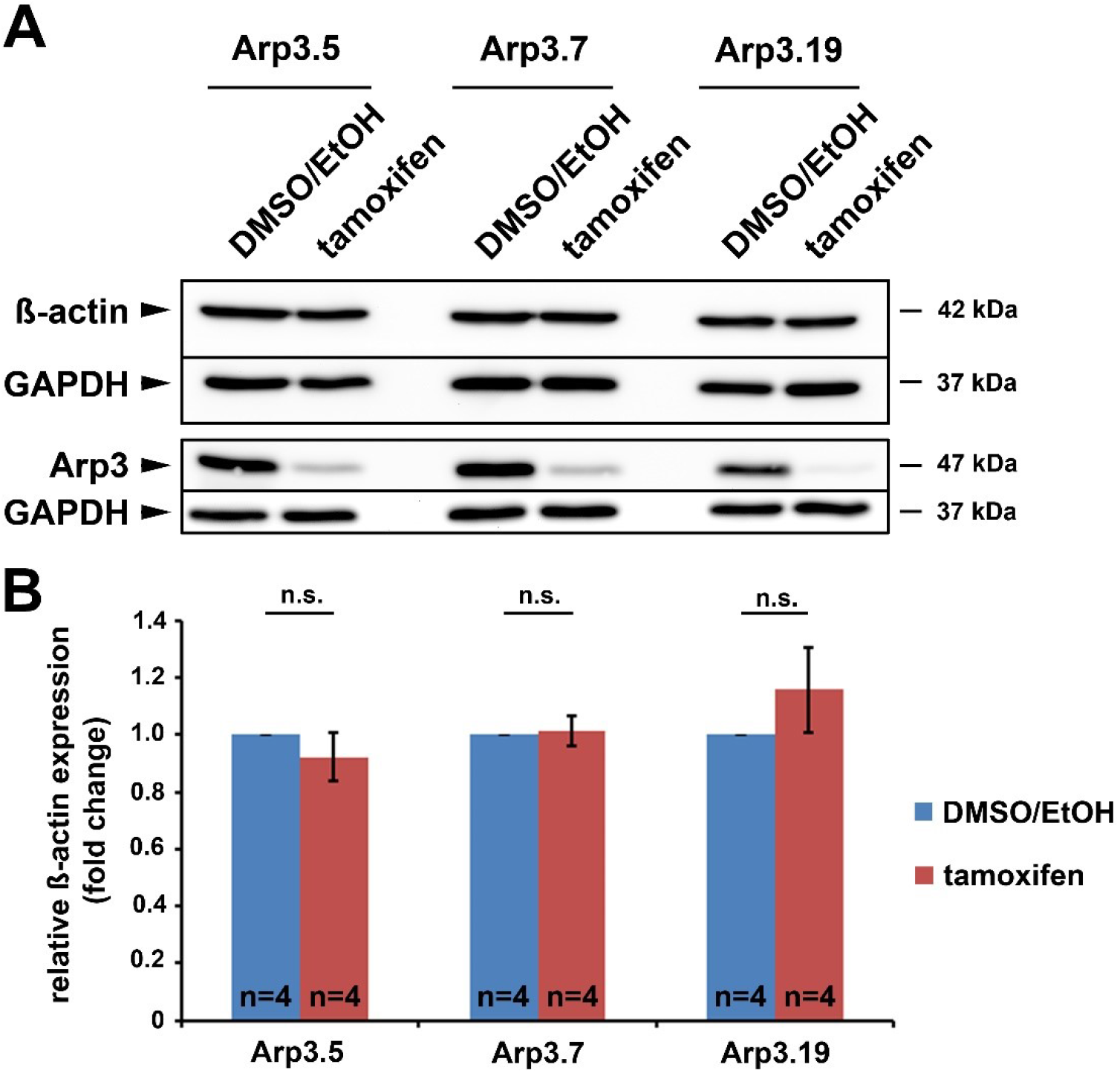
Acute Arp3 knockout does not alter ß-actin levels. **(A)** Representative Western blot of ß-actin levels in *Arp3fl/fl* clones with or without tamoxifen treatment, as indicated. GAPDH served as loading control. Note efficient Arp3 protein rundown after tamoxifen treatment. **(B)** Quantification of ß-actin levels form Western blots as shown in (A). Bar charts are arithmetic means +/− SEMs of ß-actin levels normalized to GAPDH, with each knockout (red) being presented as fold-change to its respective control (blue). Data were obtained from four independent experiments, and statistics done using non-parametric, Mann-Whitney rank sum test (n.s. not significant).

**Figure S5.**
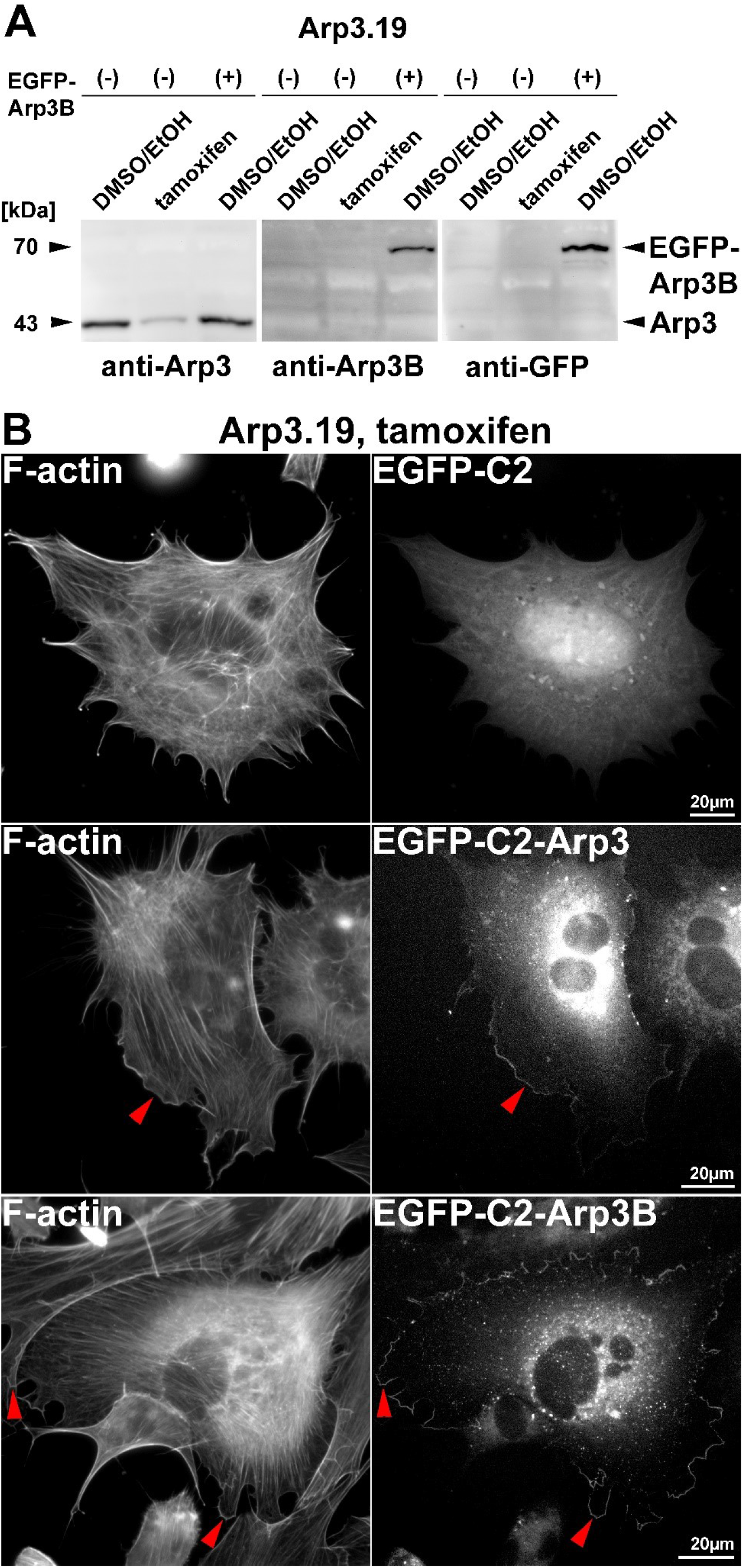
Overexpression of both Arp3 and Arp3B restores lamellipodia formation upon removal of endogenous Arp3. **(A)** Representative Western blot analysis of Arp3 and Arp3B expression in *Actr3*^*fl/fl*^ cells (clone Arp3.19) in control conditions (DMSO/EtOH) or after induction of *Actr3* knockout (tamoxifen). Arp3-specific antibody confirmed endogenous expression of Arp3 in DMSO/EtOH-treated cells as well as its successful depletion upon tamoxifen treatment. In contrast, anti-Arp3B antibody is specific for Arp3B protein, as illustrated by overexpression of EGFP-tagged Arp3B, but does not appear to be expressed (with or without Arp3 removal) from its endogenous locus (*Actr3b*). Anti-GFP antibodies served as control for EGFP-Arp3B transfection, as indicated. **(B)** Phalloidin stainings (left panels) of Arp3.19 cells following tamoxifen treatment and transient transfection with either EGFP-C2-blank, EGFP-C2-Arp3 or EGFP-C2-Arp3B (right panels), the two latter of which targeted to the cell periphery, as expected (red arrowheads). Note the restoration of lamellipodia (red arrowheads) with both EGFP-tagged Arp3 and Arp3B, but not EGFP alone, strongly suggesting the absence of expression of functional protein levels from the endogenous *Actr3b* gene upon *Actr3* disruption.

**Figure S6.**
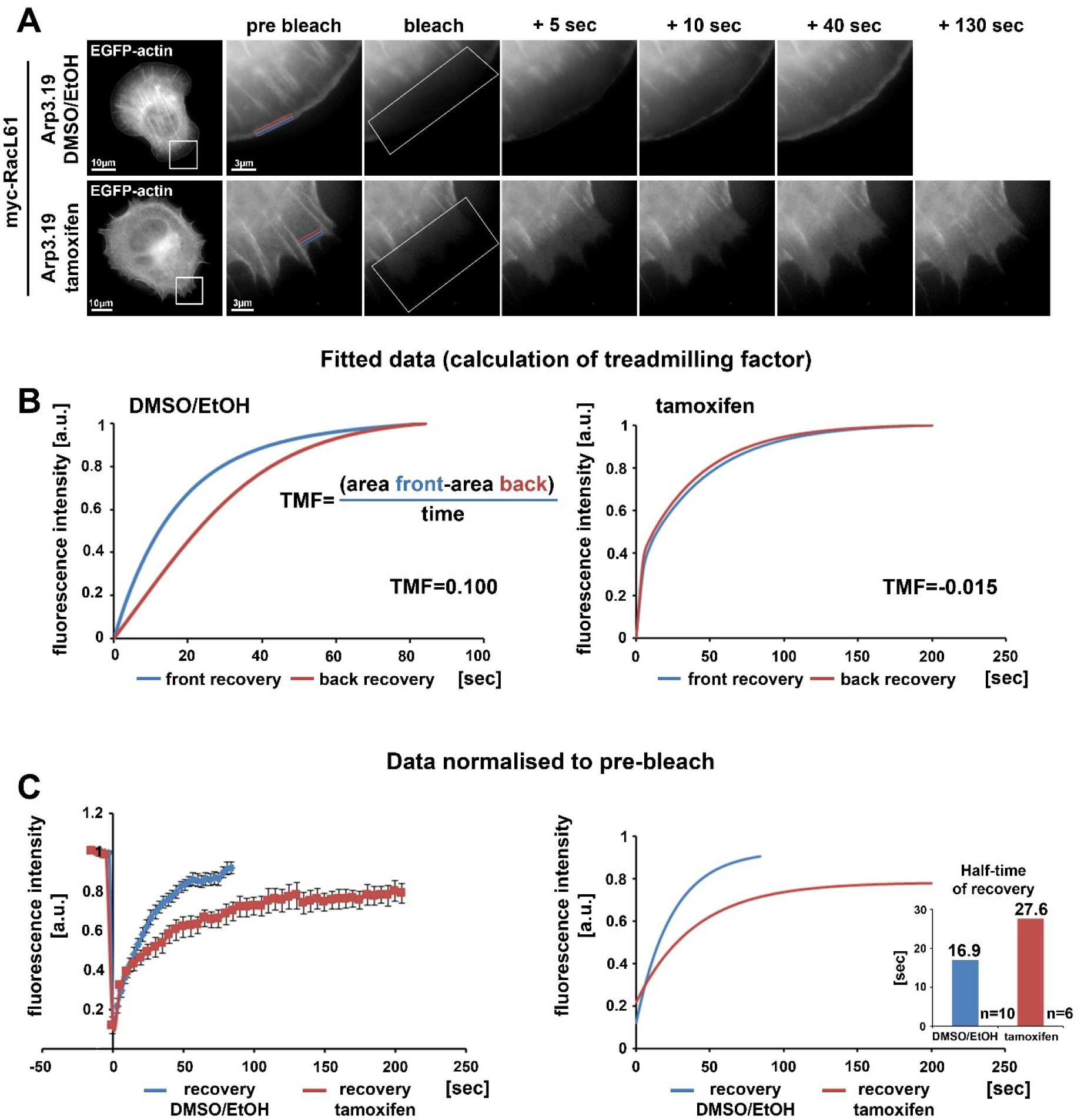
Loss of the Arp2/3 complex interferes with treadmilling and F-actin turnover even upon expression of constitutively active Rac. Experiments and analyses shown were as described for Figure 3, except that clone Arp3.19 cells were co-transfected with EGFP-actin and constitutively active Rac1. Movies from 10 control and 6 tamoxifen-treated cells acquired on three independent experimental days were analyzed.

**Figure S7.**
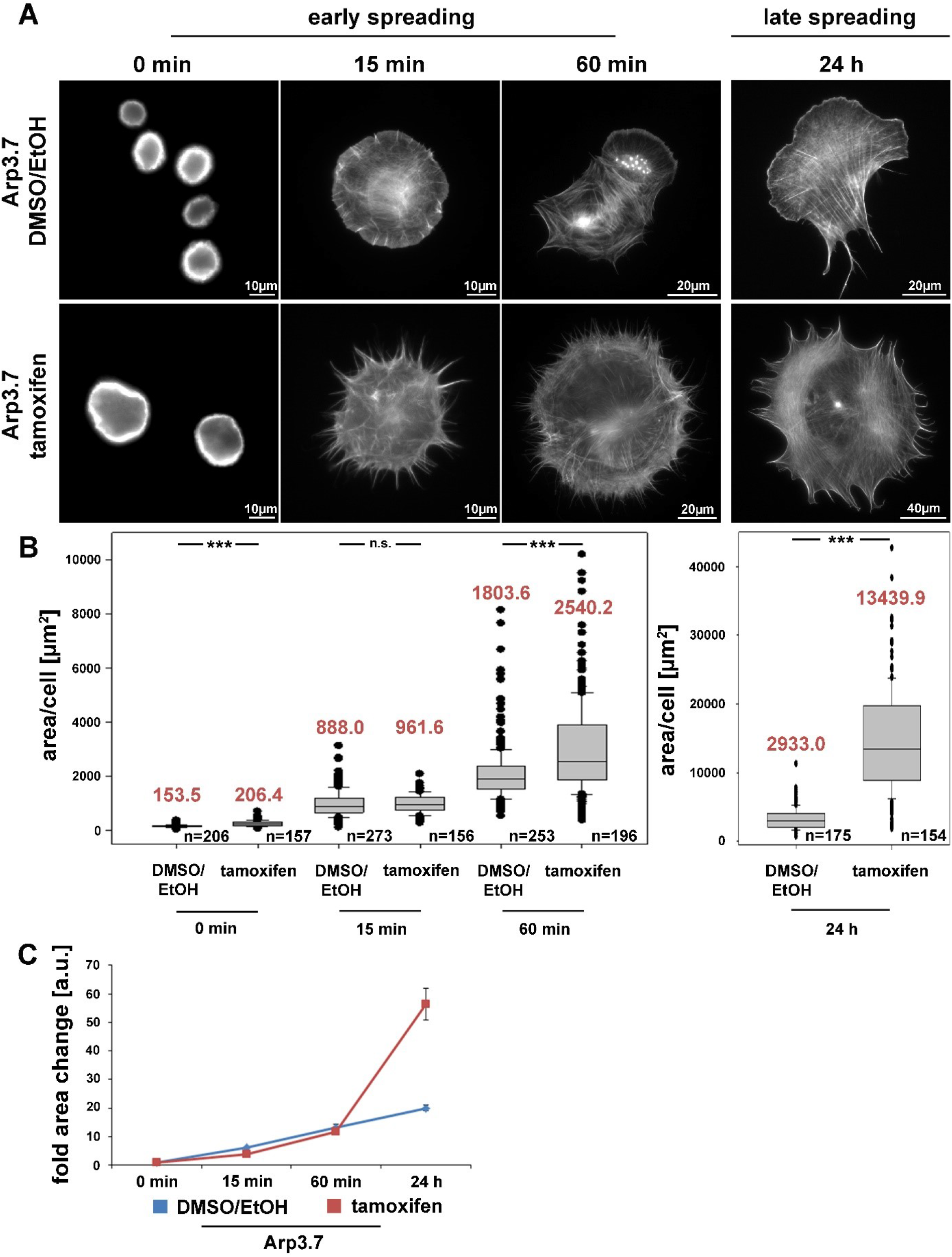
The Arp2/3 complex is not required for cell spreading, clone Arp3.7. Data generated and analyzed in analogy to those shown for clone Arp3.19 in Fig. 4.

**Figure S8.**
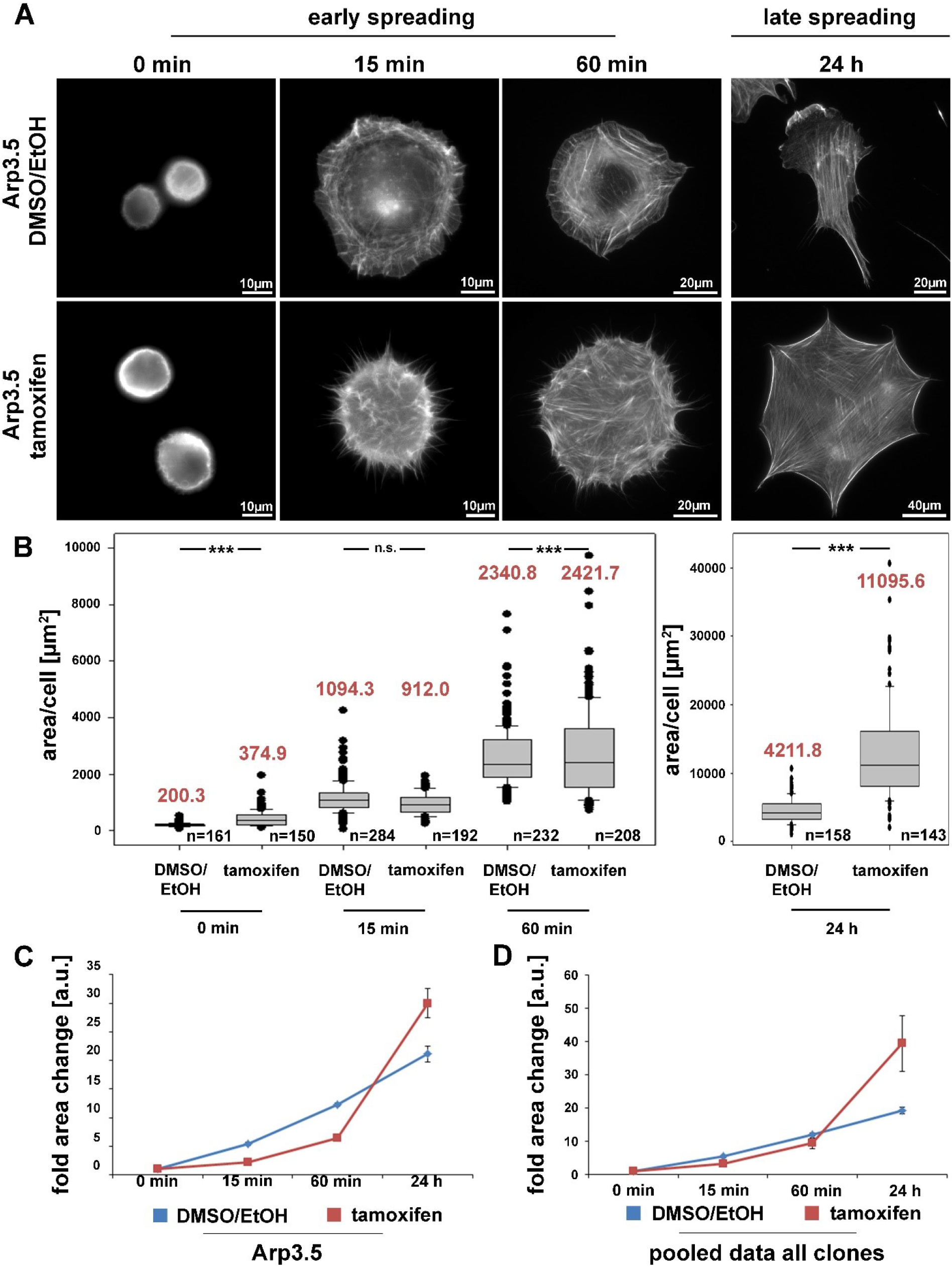
The Arp2/3 complex is not required for cell spreading, clone Arp3.5. Data generated and analyzed in analogy to those shown for clone Arp3.19 in Fig. 4. In (C), cells corresponding to clone Arp3.5 display a slight delay in spreading efficiency if normalized to the strong differences in cell size at the onset of spreading (0 min-time point), which was interpreted as clonal variation as opposed to genotype-dependent difference, because not seen for clones Arp3.19 (Fig. 4) and Arp3.7 (Fig. S7) treated with tamoxifen. This view is confirmed when pooling data from all three clones, as shown in **(D)**.

**Figure S9.**
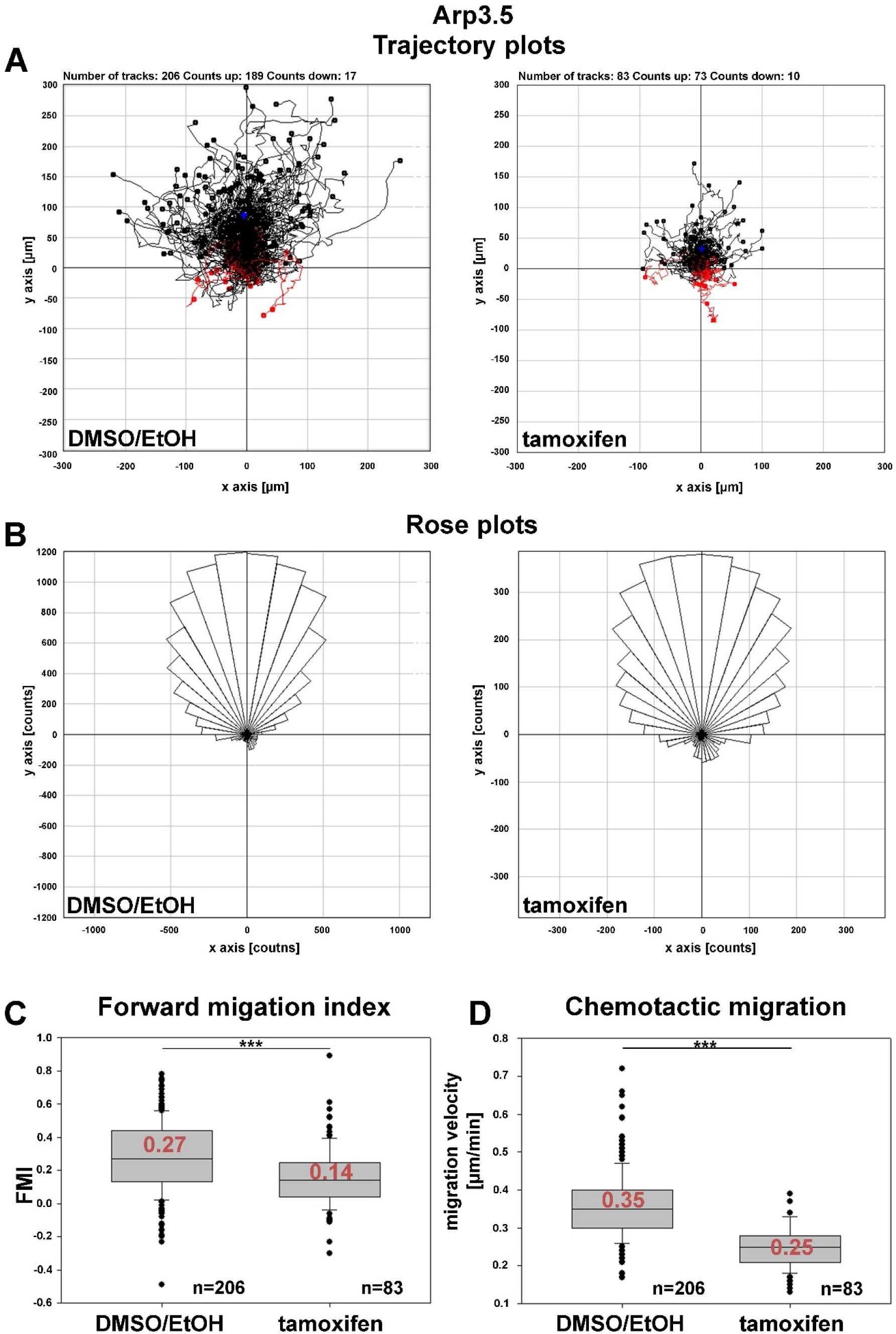
Arp2/3 complex contributes to, but is not essential for chemotaxis, clone Arp3.5. Data generated and analyzed in analogy to those shown for clone Arp3.19 in Fig. 7.

**Figure S10.**
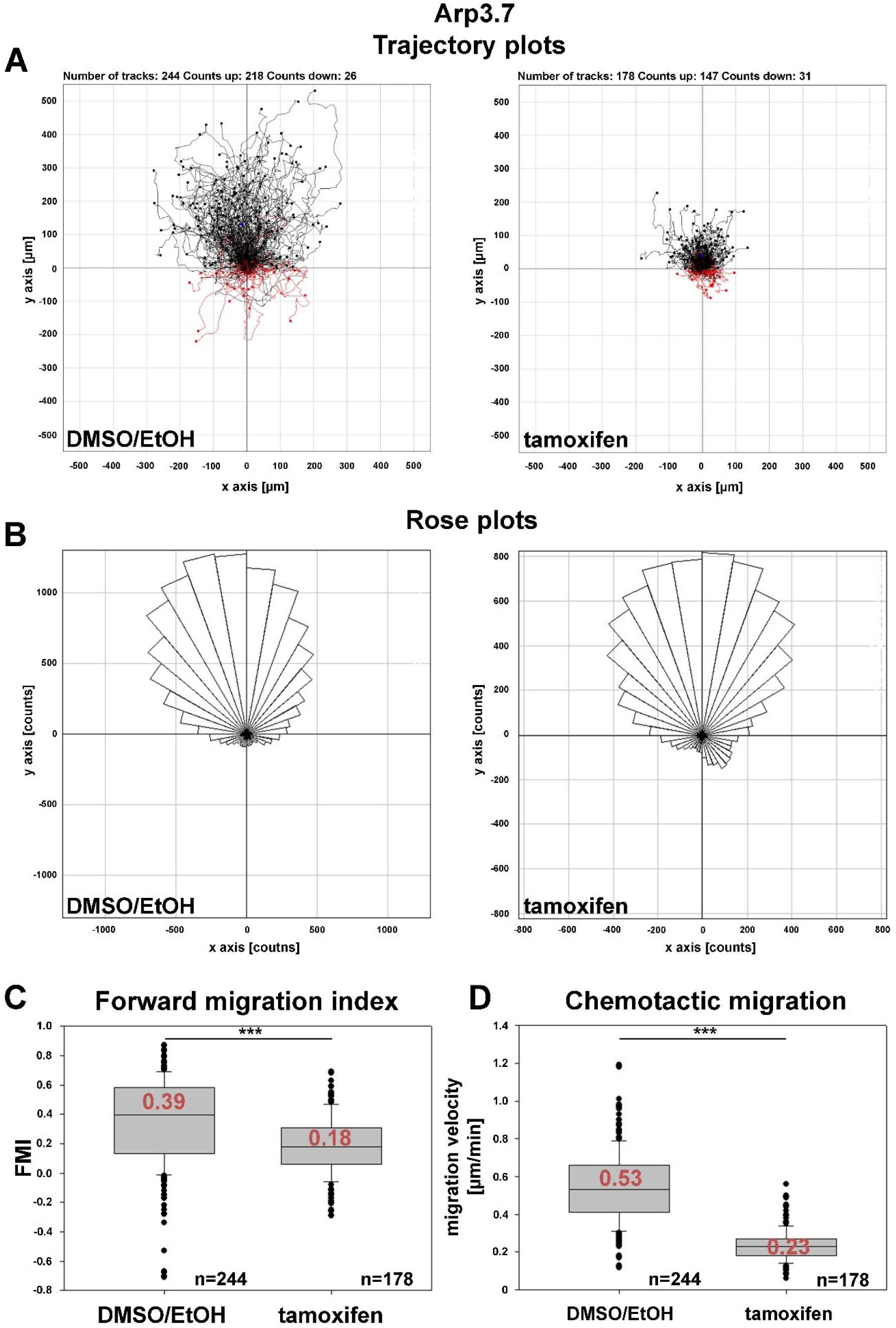
Arp2/3 complex contributes to, but is not essential for chemotaxis, clone Arp3.7. Data generated and analyzed in analogy to those shown for clone Arp3.19 in Fig. 7.

**Figure S11.**
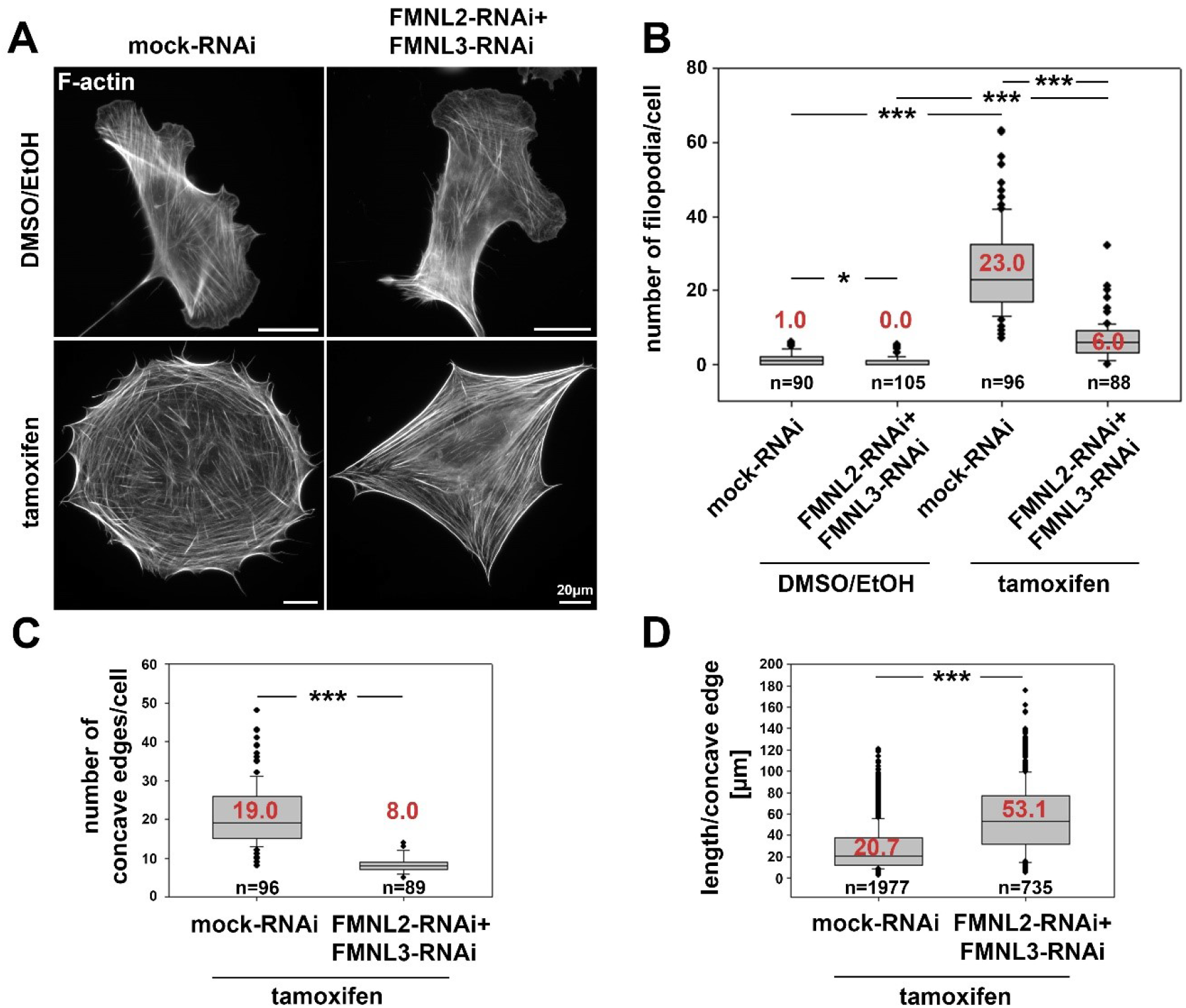
Depletion of FMNL formins reduces cell edge complexity in Arp3 knockout fibroblasts. **(A)** Representative phalloidin stainings of spread, DMSO/EtOH- and tamoxifen-treated *Actr3*^*fl/fl*^ MEFs (clone Arp3.19) subjected to either mock-RNAi or FMNL2+3-RNAi. **(B)** Quantification of filopodia numbers/cell in conditions as depicted in (A). Data are presented as box and whiskers plots as described for Fig. 4B. n equals total number of cells analyzed from three independent experiments. Differences in filopodia numbers/cell between experimental conditions were confirmed to be statistically significant using non-parametric, Mann-Whitney rank sum test (* p<0.05; *** p<0.001). Note the induction of filopodia formation by acute Arp3 removal even in readily spread cells, which appears largely effected by elevated FMNL protein levels, as quantified in Fig. 9B. **(C/D)** Box and whiskers plots displaying manual quantification of the number of concave edges defining the cell periphery of tamoxifen-treated *Arp3fl/fl* cells and length measurements of those concave edges (D) using images as displayed in (A). n corresponds to the total number of cells analyzed (C) and total number of measured concave edges (D) from three independent experiments. Statistics was done using non-parametric, Mann-Whitney rank sum test (*** p<0.001). Note that upon depletion of Arp3, the cell edge is not only characterized by an increase in filopodia numbers, but also by the formation of many concave edges of short length (A). However, the number of these concave edges is significantly reduced with the length of each concave edge being significantly increased once Arp3 removal is combined with RNAi-mediated suppression of FMNL2/3 formin expression. As a result, cells reduced for both Arp3 and FMNL2/3 expression display less complex cell edges (C) with concave edges being increased in length (D) as compared to cells lacking Arp3 alone.

**Figure S12.**
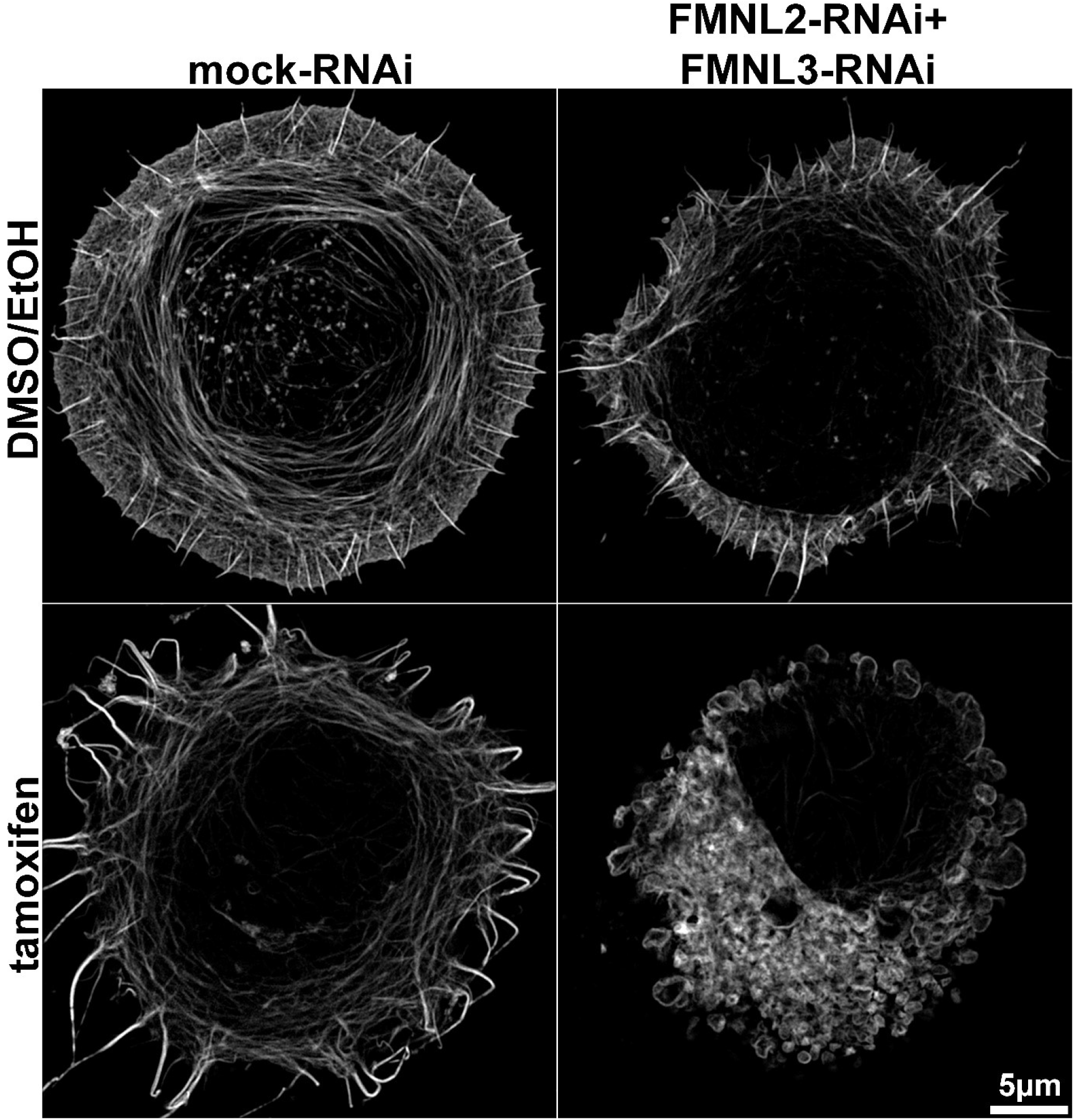
Morphological consequences of individual and combined interference with Arp3 and FMNL2/3 formin function. Structured illumination microscopy (SIM) images of phalloidin-stained cells allowed to spread for 15 minutes following genetic as well as combined RNAi treatments, as indicated.

**Figure S13.**
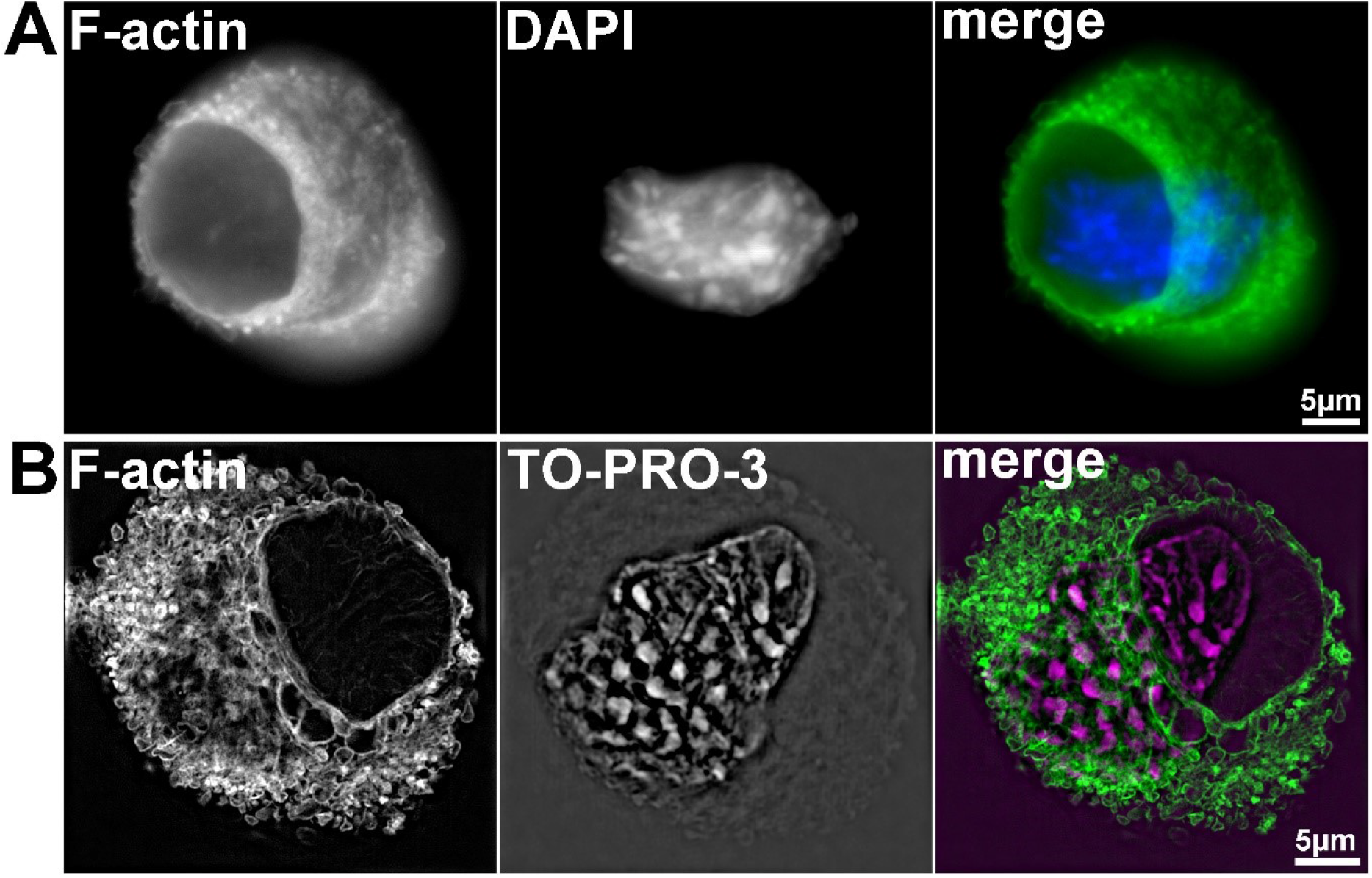
Morphology of the actin cytoskeleton in spreading cells simultaneously suppressed for Arp3 and FMNL2/3 expression. **(A)** Widefield fluorescence and **(B)** SIM imaging of Arp3 and FMNL2/3-depleted, spreading cells stained with phalloidin (F-actin) and the nucleus, as indicated. Note the formation of an adhesion area of reduced F-actin intensity, which did not simply coincide with the position of the nucleus.

**Figure S14.**
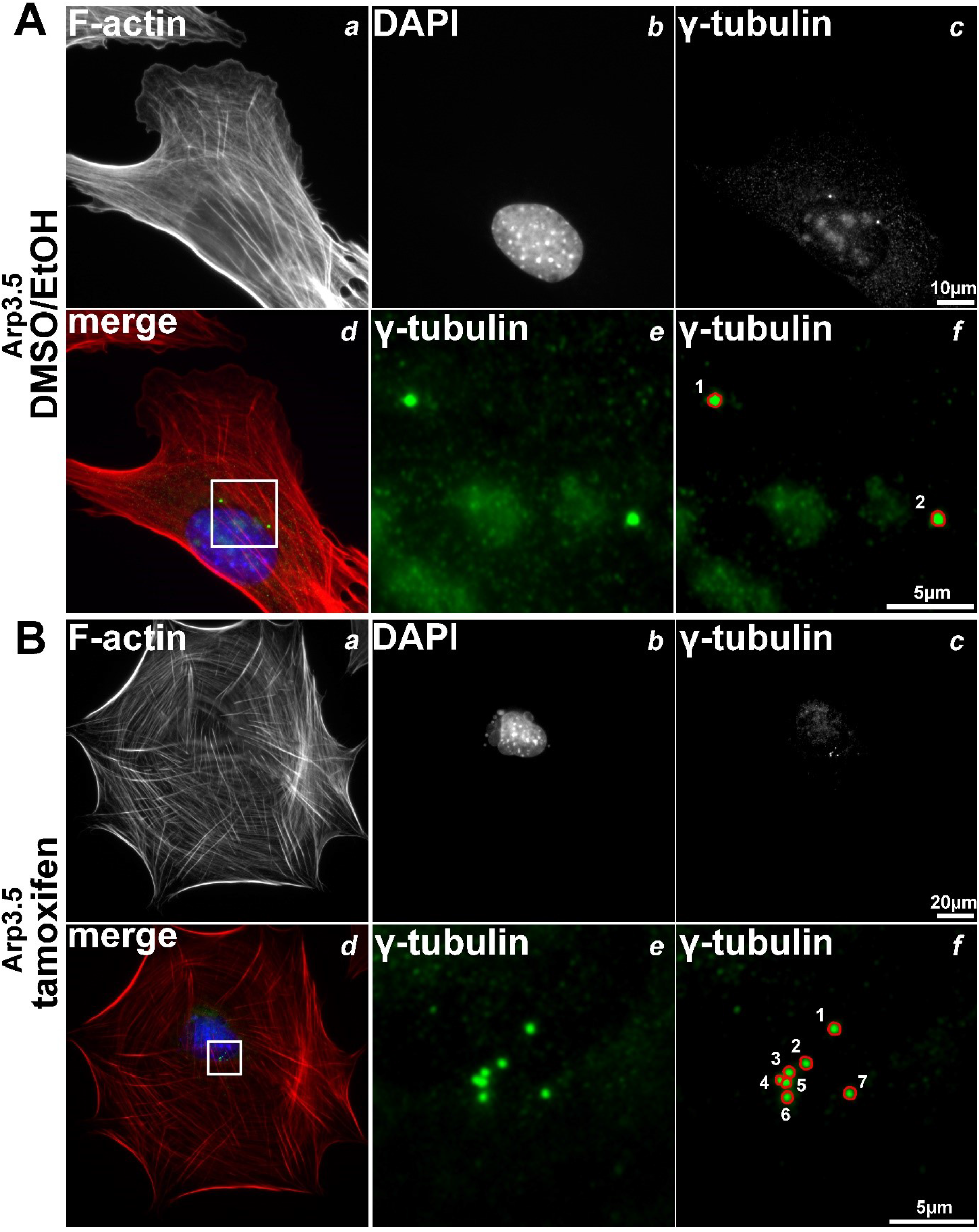
Arp3 depletion causes an increase of centrosome and nucleus numbers as well as nuclear deformation, clone Arp3.5. Representative raw data for clone Arp3.5 corresponding to quantifications shown in Fig. 10C-E.

**Figure S15.**
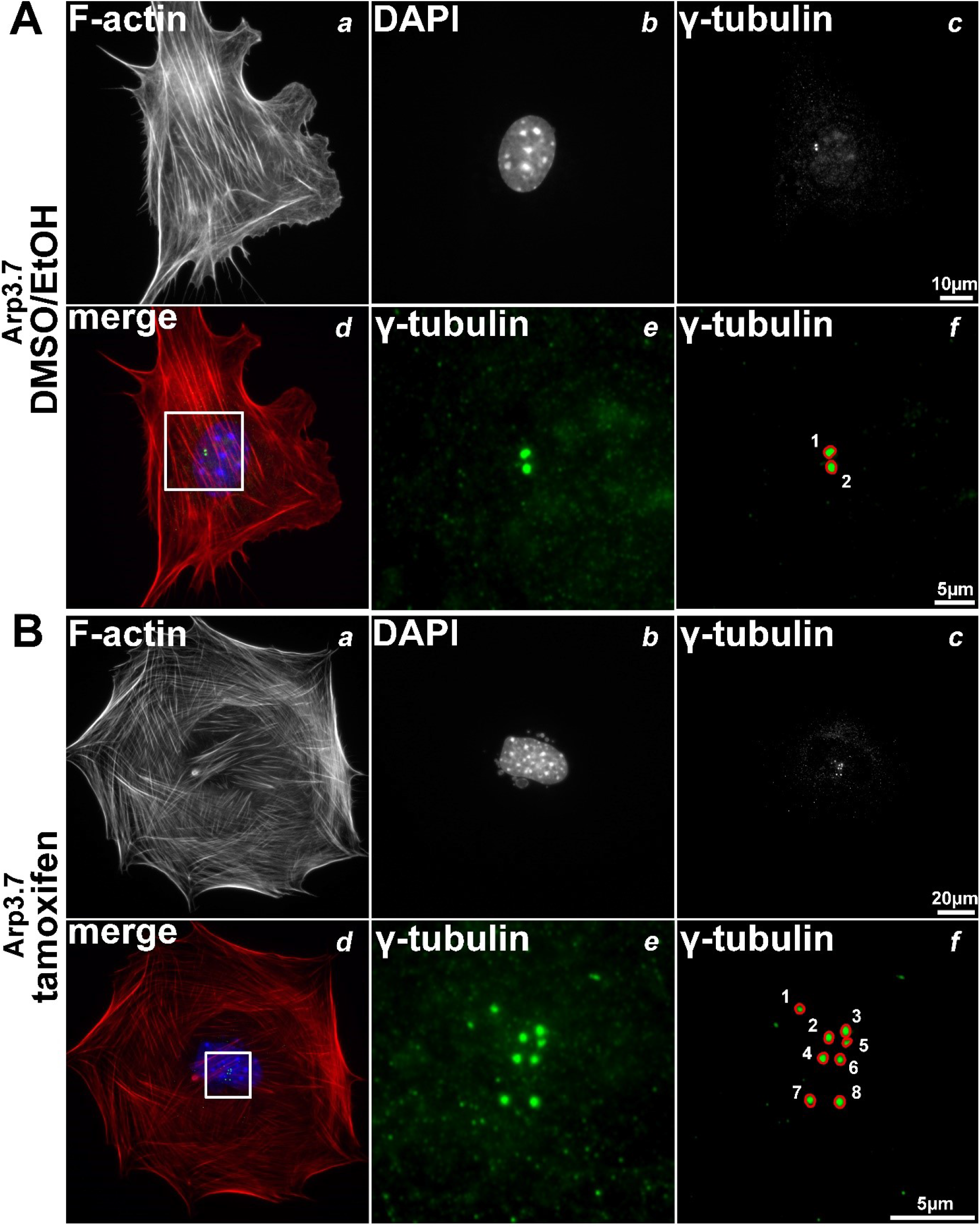
Arp3 depletion causes an increase of centrosome and nucleus numbers as well as nuclear deformation, clone Arp3.7. Representative raw data for clone Arp3.7 corresponding to quantifications shown in Fig. 10C-E.

### Legends to supplementary videos

**Video S1. Arp2/3 complex knockout fibroblasts efficiently spread using numerous filopodia.** Time-lapse microscopy of *Actr3*^*fl/fl*^ MEFs (clone Arp3.7) with (right) or without (left) tamoxifen treatment, and allowed to spread on fibronectin. Time is given in minutes and seconds.

**Video S2. Wound healing migration with and without acute reduction of Arp2/3 complex function.** Time-lapse microscopy of *Actr3*^*fl/fl*^ MEFs (clone Arp3.5) with (right) or without (left) tamoxifen treatment, and stimulated to migrate upon cell monolayer wounding. Time is given in hours and minutes (bottom right of each panel), and bars at the bottom left correspond to 50μm.

**Video S3. 3D SIM projection of phalloidin-stained control cell.** DMSO/EtOH-treated *Actr3*^*fl/fl*^ cell (clone Arp3.19) transfected with mock-RNAi vector and allowed to spread for 15 minutes.

**Video S4. 3D SIM projection of phalloidin-stained Arp3/FMNL2/3-depleted cell.** Tamoxifen-treated *Actr3*^*fl/fl*^ cell (clone Arp3.19) transfected with FMNL2+3-RNAi vectors and allowed to spread for 15 minutes. Cells suppressed for both Arp2/3 complex and FMNL2/3 formins appear to adopt a highly peculiar shape (thick flattened disks), likely caused by actin assembly deficiencies at the cell periphery caused by this treatment.

**Video S5. 3D SIM projection of phalloidin-stained Arp3/FMNL2/3-depleted cell, representative example 2.** Tamoxifen-treated *Actr3*^*fl/fl*^ cell (clone Arp3.19) transfected with FMNL2+3-RNAi vectors and allowed to spread for 15 minutes. Note that in this case, the optical sectioning allows a view from inside the cell, revealing the typical F-actin-deficient area (see also legend to Fig. S13) to be positioned at the ventral cell surface.

**Video S6. 3D SIM projection of Arp3/FMNL2/3-depleted cell co-stained with phalloidin (green) and the nucleus with TO-PRO-3 (pink).** Tamoxifen-treated *Actr3*^*fl/fl*^ cell (clone Arp3.19) transfected with FMNL2+3-RNAi vectors and allowed to spread for 15 minutes. Again, the F-actin-deficient zone at the ventral cell surface does not fully co-inside with the position of the nucleus.

